# Biosynthesis of Strained Amino Acids Through a PLP-Dependent Enzyme via Cryptic Halogenation

**DOI:** 10.1101/2023.12.13.571568

**Authors:** Max B. Sosa, Jacob T. Leeman, Lorenzo J. Washington, Henrik V. Scheller, Michelle C. Y. Chang

## Abstract

Amino acids (AAs) are modular and modifiable building blocks which nature uses to synthesize both macromolecules, such as proteins, and small molecule natural products, such as alkaloids and non-ribosomal peptides (NRPs). While the 20 main proteinogenic AAs display relatively limited side-chain diversity, a wide range of non-canonical amino acids (ncAAs) exist that are not used by the ribosome for protein synthesis but contain a broad array of structural features and functional groups not found in proteinogenic AAs. In this communication, we report the discovery of the biosynthetic pathway for a new ncAA, pazamine, which contains a cyclopropane ring formed in two steps. In the first step, a chlorine is added onto the C_4_ position of lysine by a radical halogenase PazA. The cyclopropane ring is then formed in the next step by a pyridoxal-5’-phosphate-dependent enzyme, PazB, via an S_N_2-like attack onto C_4_ to eliminate chloride. Genetic studies of this pathway in the native host, *Pseudomonas azotoformans*, show that pazamine and its succinylated derivative, pazamide, potentially inhibit ethylene biosynthesis in growing plants based on alterations in the root phenotype of *Arabidopsis thaliana* seedlings. We further show that PazB can be utilized to make an alternative cyclobutane-containing AA. These discoveries may lead to advances in biocatalytic production of specialty chemicals and agricultural biotechnology.

---

α-Amino acids (AAs) serve as a diverse group of chiral building blocks used to construct a broad range of structures in either a templated or non-templated fashion by biological systems. A subset of twenty standard AAs is genetically encoded for ribosomal protein synthesis and includes aliphatic, aromatic, acidic, and basic sidechains. However, these proteinogenic AAs represent only a small fraction of the chemical diversity found in nature.^[1]^ Indeed, both the AA monomers themselves and their resulting peptides can be highly modified to produce both new non-canonical AAs (ncAAs) and peptides of complex structure made either ribosomally^[2]^ or non-ribosomally.^[3]^ These downstream reactions include structural changes that affect backbone structure such as epimerization,^[4]^ *N*-alkylation,^[5]^ *N*-hydroxylation,^[6]^ and cyclization.^[7]^ Modification also occurs to incorporate new functionalities, such as hydroxyl groups^[8]^, halogens,^[9]^ alkenes and alkynes,^[10]^ N-N bonded motifs like diazo,^[11]^ hydrazine,^[12]^ and diazeniumdiolate groups,^[13]^ oxidized amines like hydroxylamines and nitro^[14]^ groups, heterocycles like aziridines^[15]^ and azetidines^[16]^, or incorporation of unusual elements such as fluorine,^[17]^ arsenic,^[18]^ and selenium.^[19]^ As such, AAs serve as an important source of diversity generation in biosynthesis, yielding the rich natural product structure found in ncAAs, alkaloids, nonribosomal peptides (NRPs), ribosomally-synthesized and post-translationally modified peptides (RiPPs) as well as new functionality in proteins found in post-translationally derived cofactors.^[20,21]^

In particular, the study of ncAA biosynthesis has uncovered many interesting structural motifs and biosynthetic transformations.^[1]^ After the identification of a new strategy for terminal alkyne formation in the ncAA, β-ethynylserine (βes), through cryptic chlorination^[10]^ we became interested in exploring the role of other BesD halogenases in biosynthesis of AA-derived natural products. The BesD family was the first Fe(II)/α-ketoglutarate (αKG)-dependent radical halogenase family found to halogenate free AAs by activating C(*sp*^3^)-H bonds on the methylene backbone.^[9]^ Bioinformatic analysis shows that they are located in a wide range of biosynthetic contexts and we were therefore interested to explore the different outcomes.

In this work, we describe the discovery of the cyclopropane amino acid pazamine (**1**) and its derivative pazamide (**2**) (*Figure 1A*). Pazamine is made from lysine via a remarkably efficient two step pathway consisting of PazA and PazB, where radical chlorination by PazA enables a pyridoxal phosphate (PLP)-dependent cyclopropanation carried out by PazB. Studies with *Arabidopsis thaliana* seedlings suggest that the physiological function of **1** could be related to inhibition of ethylene biosynthesis. Furthermore, PazB can produce carbocycles of different ring sizes and be applied to the biocatalytic production of a cyclobutane amino acid.

**Figure 1.**
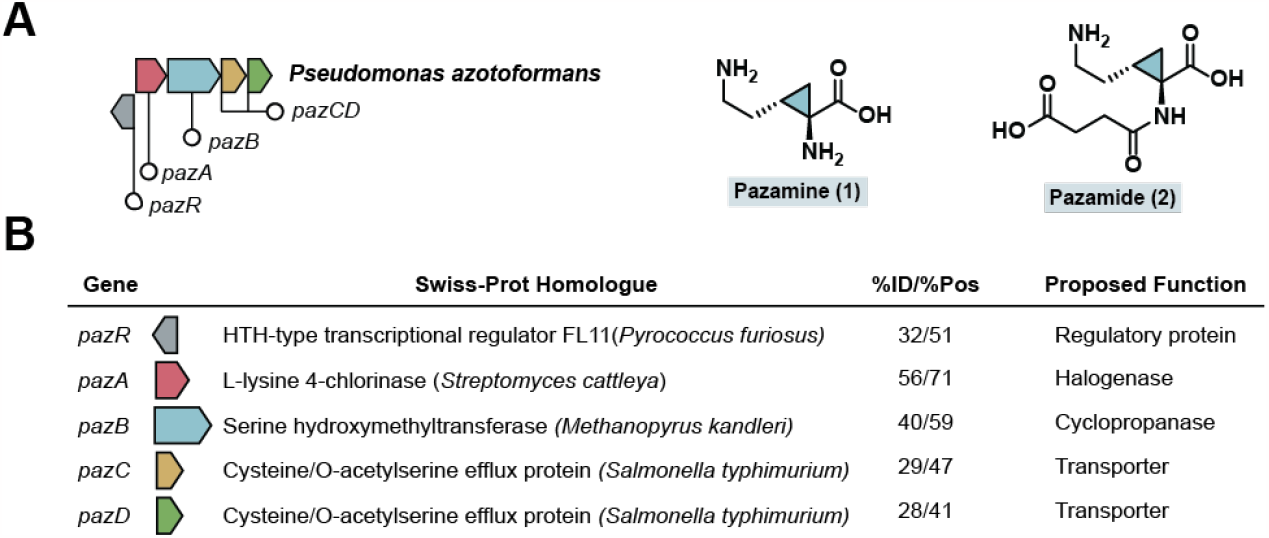
Discovery of a biosynthetic cluster that produces a cyclopropane amino acid. (A) A biosynthetic gene cluster required for the biosynthesis of cyclopropane-containing amino acid **1** and its downstream derivative **2** in *P. azotoformans*. (B) A bioinformatic analysis of the genes in the *pazRABCD* gene cluster. The genes are annotated based on Swiss-Prot homologues identified by a BLAST search and the percent identity (%ID) and percent similarity (%Pos) are shown. The proposed functions of the genes are included, based on the results of this study.

The *pazRABCD*, cluster was first identified in *Pseudomonas azotoformans* through its BesD-like halogenase. The cluster encodes a regulatory protein (PazR), a lysine halogenase (PazA), a serine hydroxymethyltransferase-like enzyme (PazB), and two putative amino acid transporters (PazCD) (*Figure 1B, Table S1, Figure S1*). Serine hydroxymethyltransferases are a class of PLP-dependent enzymes which catalyze the interconversion of serine and glycine using tetrahydrofolate (THF) as a co-substrate, as well as folate-independent aldolase chemistry (*Figure S2)*.^[22]^ A bioinformatic analysis revealed that PazB is distinct from the canonical bacterial SHMTs and a majority of its homologs are found in archaea (*Figures S1-S2*). Furthermore, residues involved in folate-dependent chemistry are not present in PazB, even though they are highly conserved in SHMTs from all domains of life (*Figure S2*).^[23,24]^ Coupled with the fact that *pazB* is genetically colocalized with an amino acid halogenase, this information suggests that PazB may carry out a novel function related to the modification of (2*S*,4*R*)-chlorolysine produced by PazA. Additionally, this cluster and larger *pazAB*-containing clusters are found in other *Pseudomonas* spp. while select *Legionella* spp. encode a similar cluster containing a C_4_-lysine dichlorinase homolog (*Figure S3*).

We set out to characterize the product of the *pazRABCD* gene cluster. Since PazB expresses very little soluble protein in *Escherichia coli* (*Table S2*), these studies were carried out in the native host, *P. azotoformans*. The PazAB overexpression plasmid, pPazAB, was constructed by inserting the native *pazAB* fragment into the pMMPc-Gm plasmid under control of the Pc promoter from *Delftia acidovorans*.^[25]^ The *P. azotoformans* pPazAB overexpression strain was then cultured in parallel with the *P. azotoformans* pMMPc-Gm empty plasmid control strain. The intracellular metabolome was extracted with acidic methanol and analyzed by high-resolution liquid chromatography-quadrupole time-of-flight mass spectrometry (LC-QTOF). Differences in the metabolomes of these two strains were assessed using MS-DIAL^[26]^ (*Figure S4*), revealing a metabolite (*m*/*z* = 245.1132 [M+H]^+^) that is overexpressed in *P. azotoformans* pPazAB but absent in the control strain (*Figure 2A*). Based on the calculated molecular formula (C_10_H_16_N_2_O_5_) and the absence of a ^37^Cl isotope pattern in the mass spectrum, it seemed possible that the biosynthesis of the PazAB product could involve cryptic chlorination. *Pseudomonas azotoformans* pPazAB was grown in the presence of the isotopically labeled precursors 4,4’,5,5’-*d*_4_-L-lysine and ^13^C_6_,^15^N_2_-L-lysine to further validate this metabolite’s origins. These studies showed that L-lysine is indeed incorporated into this product and that a deuteron from either C_4_ or C_5_ of lysine is removed, consistent with radical halogenation of lysine by PazA. In addition to lysine, there was another four-carbon fragment incorporated into the product. This fragment was determined to originate from succinate, which was confirmed by feeding experiments with *d*_4_-succinate (*Figure 2B*).

**Figure 2.**
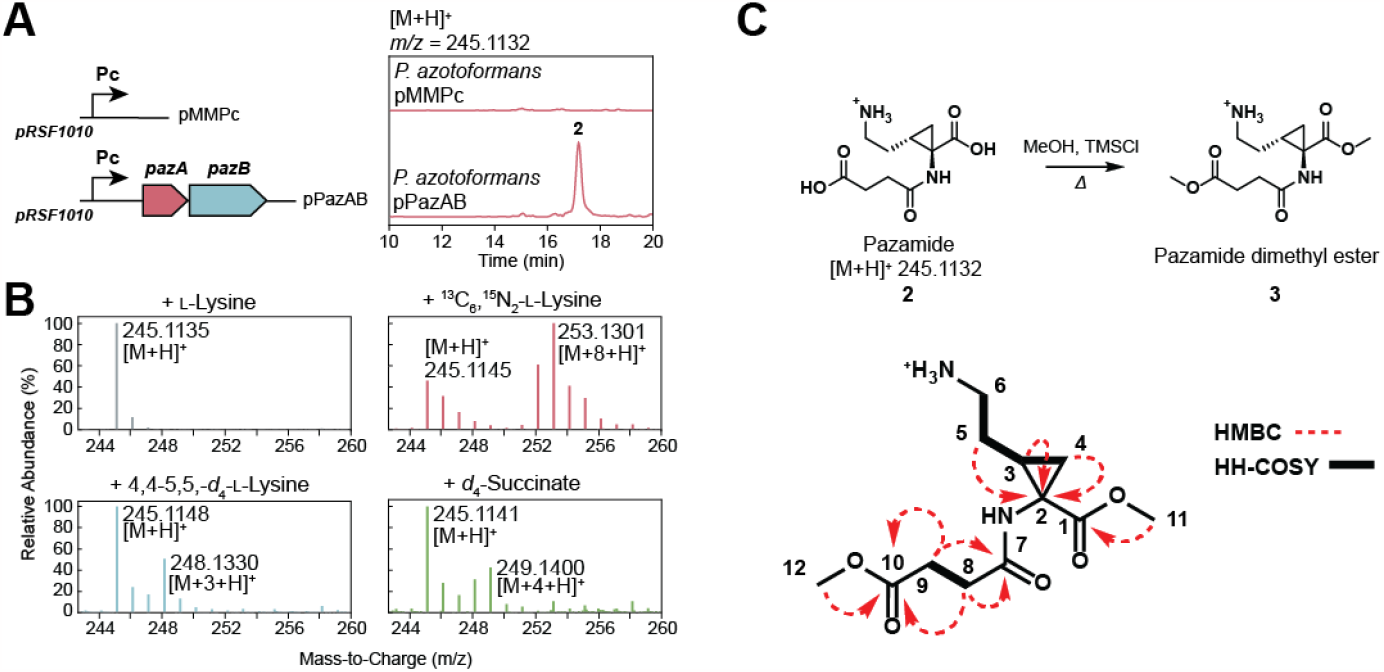
PazAB produces a cyclopropane amino acid (A) *P. azotoformans* transformed with either the PazAB expression plasmid (pPazAB) or the empty vector control (pMMPc) were cultured. Metabolomic analysis of the extracted cell culture revealed a new metabolite of *m/z* = 245.1132 [M+H]^+^ that was only detected when PazAB were expressed. (B) Isotopic labeling studies showed that the M+8 isotopologue is formed upon feeding of ^13^C_6_,^15^N_2_-L-lysine, confirming that the product is derived from L-lysine. Production of the M+3 isotopologue upon feeding of 4,4’,5,5’-*d*_4_-L-lysine is consistent an H-abstraction step catalyzed by PazA during chlorination. Feeding of *d*_4_-succinate leads to production of the M + 4 isotopologue, indicating that succinate is the source of the additional four carbon fragment . (C) Compound **2** was isolated and methyl esterified for structural elucidation by 2D-NMR techniques. Key HMBC correlations are shown as red arrows while HH-COSY correlations are indicated with bold lines.

This compound was purified by preparative HPLC using hydrophilic interaction chromatography (HILIC). The product was analyzed by 2D-NMR following methyl esterification and determined to be a succinylated cyclopropane-containing amino acid (**3**) (*Figure 2C, Figure S5-S7*). The free amino acid parent compound **1** and its succinylated derivative **2** have been given the trivial names pazamine and pazamide, respectively, after the host organism *P. azotoformans*. With the product structure in hand, a biosynthetic hypothesis for **1** can be proposed where the PazA halogenase initiates the pathway by radical chlorination of the C_4_ of L-lysine (**4**) (*Figure 3A*). *In vitro* reactions of PazA with L-lysine confirm that it does indeed produce (2*S*,4*R*)-chlorolysine (**5**) (*Figure S8*), which serves as a substrate for PazB. PazB is a PLP-dependent enzyme, suggesting that the carbocycle is made via deprotonation of C_α_ followed by nucleophilic attack on C_4_ with the chloride serving as a leaving group.

**Figure 3.**
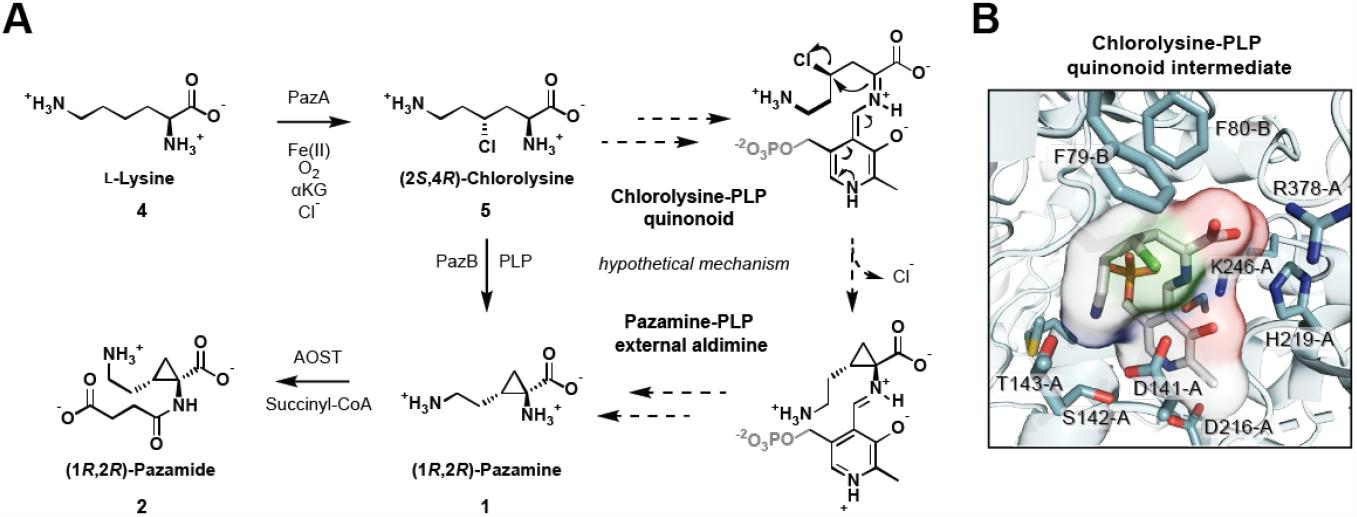
Proposed biosynthetic pathway to pazamine and pazamide (A) Cyclopropanation is proposed to take place in two steps. In the first step, radical halogenation of **4** by PazA incorporates a leaving group for the next step. In the second step, the PLP-dependent enzyme PazB reacts with **5** to form the external aldimine. C_α_-deprotonation yields the quinonoid carbanion intermediate that can carry out an S_N_2-like reaction at C_4_ to form the carbocycle with stereoinversion and concomitant loss of chloride. **1** is then released from the PLP cofactor and is succinylated by AOST to produce **2**. (B) A model of the PazB active site was generated by AlphaFold and the quinonoid intermediate before carbocyclization was docked in with AutoDock Vina. In this model, a polar pocket is observed and proposed to bind the ε-ammonium group in order to bring C_4_ into proximity of C_α_ for cyclization. The bending of the chain also appears to be assisted by steric interactions with F79 and F80.

Several PazB orthologs were tested but found to be insoluble for heterologous expression in *E. coli*. (*Table S2*). Thus, PazB was modeled using AlphaFold^[27,28]^ in order to gain more insight into the cyclopropanation reaction (*Figure S9*). AutoDock Vina^[29,30]^ was then used to generate docked poses of the putative quinonoid intermediate of PLP and **5**. Comparison to the crystal structure of a canonical SHMT from *E. coli* bound to substrate^[23]^ (PDB: 1DFO) assisted in identifying a biologically relevant pose (*Figure 3B*). Analysis of the model revealed the presence of a polar pocket in the PazB active site (D141-A, T142-A, and S143-A) that may interact with the N_ε_ of PLP-bound **5** (*Figures 3B and S9B)*. This interaction appears to facilitate folding of the methylene chain into a reactive conformation so that C_4_ can be brought into proximity of the C_α_ carbanion nucleophile. Additional steric interactions with F79-B and F80-B further appear to control positioning of the aminoalkyl chain into a productive conformation for cyclopropanation. Intramolecular attack by the C_α_ carbanion nucleophile on the chlorinated C_4_ results in chloride elimination and C-C bond formation with stereoinversion to yield the cyclopropane ring of **1**.

Given the absence of a gene candidate for a succinyltransferase in the *pazRABCD* operon or its genome neighborhood, we hypothesized that formation of **2** from **1** was not carried out by a dedicated enzyme but by an enzyme from primary metabolism. Examination of metabolic pathways involving the structurally-similar amino acid L-ornithine identified arginine/ornithine succinyltransferase (AOST), an enzyme which catalyzes the α-amino succinylation of both L-arginine and L-ornithine (*Figure S10*).^[31]^ To test this hypothesis, an AOST-knockout strain *P. azotoformans ΔaruFG* was generated. Overexpression of PazAB in this strain shows that it still produces **1** but **2** is no longer observed (*Figure S9*), which is consistent with a model where succinylation of **1** is carried out by AOST. Interestingly, AOST from *P. aeruginosa* PAO1 has been shown to be inhibited by and inactive towards D-ornithine, suggesting that **1** and L-ornithine have the same relative stereochemistry at C_α_.^[31]^ This information, the predicted stereoinversion of the cryptically chlorinated C_4_, and an NOE supported by modelling (*Figure S6*) leads to our tentative assignment of the stereochemistry of **1** (and **2**) as (1*R*, 2*R*).

As many Pseudomonads, including *P. azotoformans*, are known to associate with plant hosts,^[32]^ we thought to question the bioactivity of **1**. We noticed that **1** bears a resemblance to another cyclopropane amino acid, 1-aminocyclopropane-1-carboxylate (ACC), an intermediate in the biosynthesis of ethylene in plants. This molecule is made through the PLP-catalyzed cyclization of the methionine moiety of *S*-adenosylmethionine (SAM), liberating methylthioadenosine in the process.^[33,34]^ We thus hypothesized that **1** might interact with the ethylene pathway in a plant host, possibly as an inhibitor of either ACC synthase or ACC oxidase, lowering the amount of ethylene produced by plants *(Figure 4A)*. Plants respond to the initial detection of bacteria by inducing ethylene production prior to downstream determination of bacterial lifestyle — i.e. type of pathogen or mutualist — and subsequent immune responses.^[35]^ This results in an initial reduction of root length and increase in root branching and root hair formation unless the bacteria has processes that interfere with this response.^[36,37]^ Inoculation of *Arabidopsis thaliana* with *P. azotoformans ΔaruFG* pPazAB demonstrated a rescuing of root length inhibition caused by bacterial inoculation and subsequent ethylene production. Wild type *P. azotoformans* and the **1**-deficient *P. azotoformans ΔpazA* are unable to rescue the reduction in root length. Interestingly, *P. azotoformans* pPazAB is also unable to fully rescue the observed root phenotype, possibly because it can sequester the proposed bioactive **1** as **2** (*Figure 4B, Figure S11)*. Overall, these data align with our understanding of the plant response to bacterial inoculation. When compared to an uninoculated control, the strain altered to increase **1** production induced the expected change in root architecture, without a reduction in root length. While these results support our initial hypothesis that **1** is capable of inhibiting ethylene biosynthesis, further work is required to definitively assess if and how this occurs.

**Figure 4.**
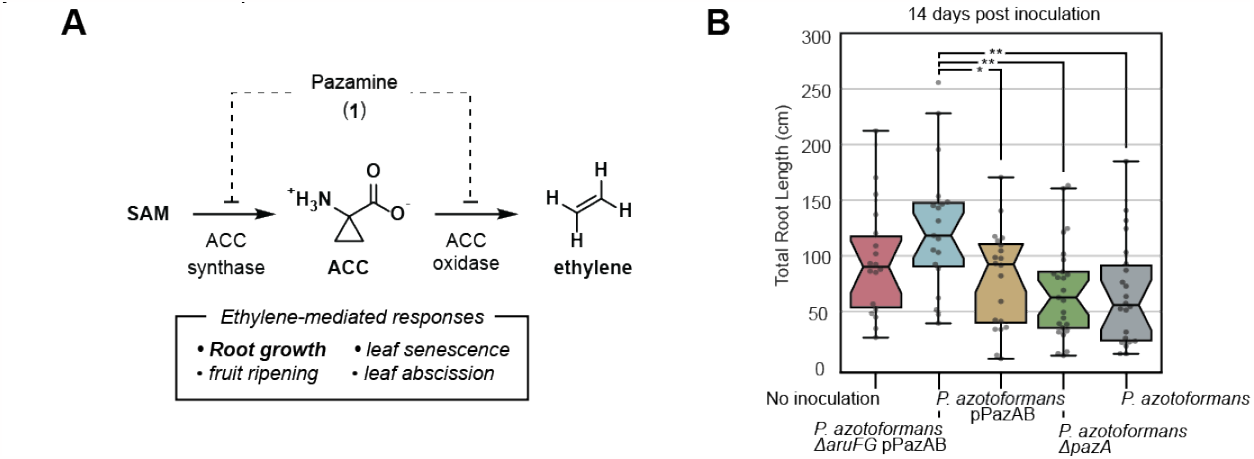
Pazamine-producing bacteria rescue root length suppression in inoculated *Arabidopsis thaliana*. (A) As **1** is structurally similar to the ethylene precursor ACC, it may be capable of inhibiting enzymes involved in plant ethylene biosynthesis. (B) Inoculation of *A*. t*haliana* with *P. azotoformans ΔaruFG* pPazAB, a **1** producer, results in a rescue of the root length phenotype caused by bacterial inoculation and subsequent ethylene production. In *P. azotoformans* pPazAB, **1** can be converted to **2**, diminishing the observed effect. This effect is statistically significantly (ANOVA and Tukey post-hoc test, * = *p*-value < 0.05, ** = *p*-value < 0.01) at 14 days post inoculation (DPI).

Strained carbocycles like the cyclopropane found in **1** are important structural elements found in a broad range of natural product structures.^[38]^ Cyclopropanes themselves are metabolically stable motifs that are present in a number of pharmaceuticals based on their rigid, biplanar nature that can enforce desired conformations while providing unique hybridization in between *sp*^2^ and *sp*^*3*^.^[39]^ Along with cyclobutane rings, they are of high interest as substrates for C-C activation and reactivity towards nucleophiles and electrophiles due to their inherent ring strain.^[40]^ In addition to synthetic applications, cyclopropane rings are important in nature in both natural products and lipids, where they they arise mostly via SAM-dependent methyl transfer to double bonds^[41]^ as well as terpene cyclase-^[42]^ and metalloenzyme-catalyzed rearrangements.^[14,43]^ Another cyclopropane amino acid from *Pseudomonas* spp., coronamic acid, is made via cryptic chlorination of a carrier protein-bound amino acid followed cyclization by a Zn^2+^-dependent enzyme.^[44]^ A handful of cyclobutane amino acids have also been found, such as 2,4-methanoproline and 2,4,-methanoglutamic acid^[45]^ but their biosyntheses are not known. Other cyclobutane-containing natural products are thought to arise from terpene cyclase-catalyzed rearrangements of terpenes or by photochemical cycloadditions.^[46]^

Considering the scarcity of biochemical methods to generate cyclobutanes, we wanted to test if PazB could accept an alternative substrate to catalyze cyclobutanation. The enzyme HalB catalyzes the stereoselective halogenation of lysine at C_5_ so co-expression of HalB with PazB could enable the production of the cyclobutane amino acid **7** (*Figure 5A*). Towards this end, we made the plasmid pJTL1 to construct this pathway and we expressed it in *P. azotoformans*. Gratifyingly, we observed the production of a unique metabolite corresponding to **7** (*m/z* = 145.092 [M+H]^+^) (*Figure 5B*). Feeding 2,3,3’,4,4’,5,5’,6,6’-*d*_9_-L-lysine to *P. azotoformans* pPazAB or pJTL1 showed that the expected M+7 isotopologues of **1** and **7** are produced, indicating that two deuterons are lost as would be consistent with radical halogenation and PazB-catalyzed cyclization (*Figure S12*). These studies show that PazB is sufficiently promiscuous to expand ring size and accept substrates that are chlorinated at either C_4_ or C_5_. Furthermore, this system provides an avenue to study enzymatic cyclobutane synthesis, a rare and underrepresented biochemical reaction, as well as a potential biocatalytic route to various carbocyclic ncAAs.

**Figure 5.**
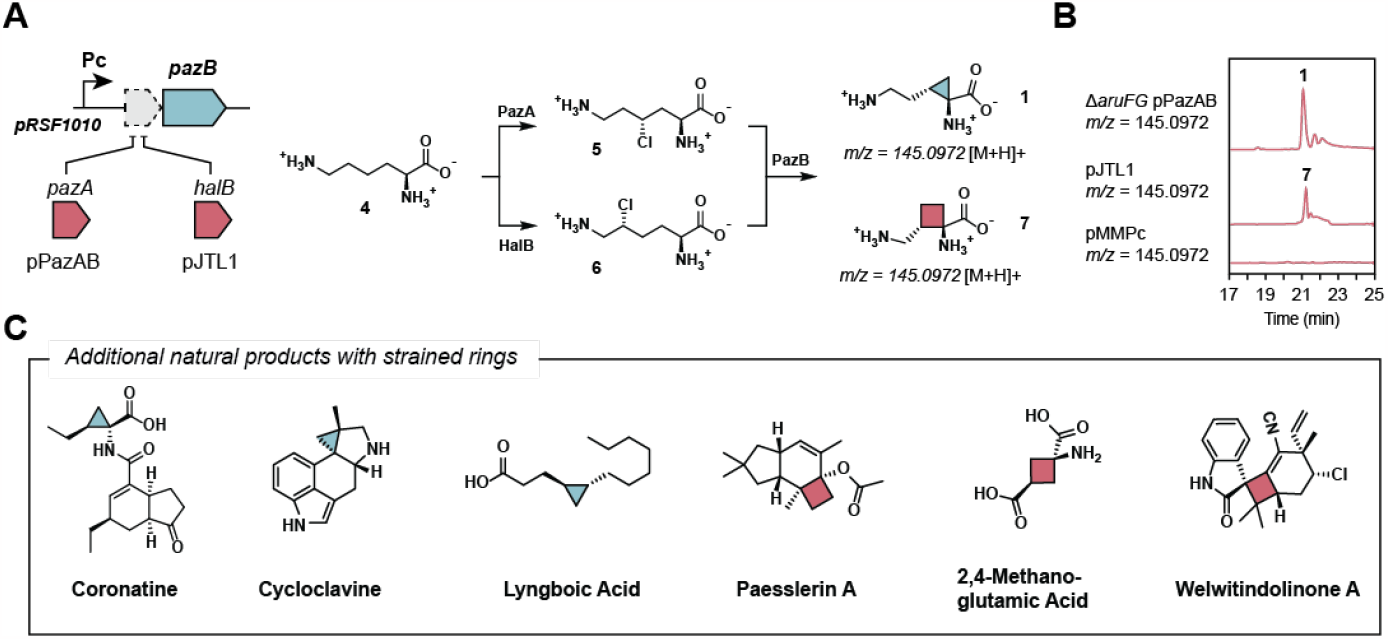
Engineering the biosynthetic production of new carbocycle-containing amino acids (A) Combinatorial expression of PazB with different halogenases allows for the biosynthesis of various strained ncAAs. HalB halogenates L-lysine (**4**) at C_5_ altering the ring size of the PazB product. (B) *P. azotoformans ΔaruFG* pPazAB and *P. azotoformans* pJTL1 cultures produce **1** and **7**, respectively. (C). Selected natural products which contain either a cyclopropane or a cyclobutane ring.

The discovery of the PazAB pathway provides an efficient enzymatic route towards formation of strained carbocyclic amino acids. The strategy of cryptic chlorination followed by PLP-catalyzed cyclization is a new variant for enzymatic cyclopropane formation that we have shown to be generalizable to production of new cyclopropane and cyclobutane amino acids. Specifically, PazB demonstrates catalytic plasticity in accepting different chlorinated amino acids for carbocycle formation, allowing ring size to be altered Analysis of the active site suggests simple electrostatic and steric interactions assist in folding the chain for cyclization, providing a design template for engineering other PLP-dependent enzymes for carbocycle formation. Indeed, the use of free amino acids by the PazAB pathway rather than tethered or specialized amino acids potentially allows greater scope for engineering new pathway variants or their incorporation into synthetic compounds or natural products utilizing amino acid building blocks. Taken together, the continual exploration of biosynthetic pathways can enable advances in biocatalysis and biosynthesis by the discovery of new reaction mechanisms and natural products for expanding the scope of enzymatic chemistry.

## Acknowledgements

This work was funded by generous support from the NIH (R01 GM134271). M.B.S. acknowledges the support of an NIH NRSA Training Grant (1 T32 GMO66698). L.J.W. acknowledges support from the NSF Graduate Research Fellowship Program (DGE 2146752). L.J.W. and H.V.S. were funded by The Novo Nordisk Foundation grant no. NNF19SA0059362 (InRoot) and by the Joint BioEnergy Institute (http://www.jbei.org) supported by the U. S. Department of Energy, Office of Science, Office of Biological and Environmental Research, through contract DE-AC02-05CH11231 between Lawrence Berkeley National Laboratory and the U.S. Department of Energy. We would also like to thank Dr. Ioannis Kiporous for generating DFT models of pazamide and Dr. Edward Koleski for generating the *P. azotoformans* ΔpazA knockout strain. Instruments in the UC Berkeley College of Chemistry NMR Facility were supported in part by NIH S10OD024998. Support for the 900 MHz NMR spectrometer in the QB3 Institute in Stanley Hall at University of California, Berkeley, was kindly provided by the NIH (GM68933).

## Supporting Information

Supporting information includes methods, sequences, along with supporting data, figures, tables, spectra, and discussion.

