## Supplemental Information for "Biosynthesis of Strained Amino Acids Through a PLP-Dependent Enzyme via Cryptic Halogenation"

**Abstract:** Amino acids (AAs) are modular and modifiable building blocks which nature uses to synthesize both macromolecules, such as proteins, and small molecule natural products, such as alkaloids and non-ribosomal peptides (NRPs). While the 20 main proteinogenic AAs display relatively limited side-chain diversity, a wide range of non-canonical amino acids (ncAAs) exist that are not used by the ribosome for protein synthesis but contain a broad array of structural features and functional groups not found in proteinogenic AAs. In this communication, we report the discovery of the biosynthetic pathway for a new ncAA, pazamine, which contains a cyclopropane ring formed in two steps. In the first step, chlorine is added onto the $C_4$ position of lysine by a radical halogenase PazA. The cyclopropane ring is then formed in the next step by a pyridoxal-5´-phosphate-dependent enzyme, PazB, via an $S_N2$-like attack onto $C_4$ to eliminate chloride. Genetic studies of this pathway in the native host, *Pseudomonas azotoformans*, show that pazamine and its succinylated derivative, pazamide, potentially inhibit ethylene biosynthesis in growing plants based on alterations in the root phenotype of *Arabidopsis thaliana* seedlings. We further show that PazB can be utilized to make an alternative cyclobutane-containing AA. These discoveries may lead to advances in biocatalytic production of specialty chemicals and agricultural biotechnology.

### Table of Contents

#### Materials and Methods


*Commercial Materials* S3
*Bacterial Strains and Culture* S3
*Bacterial Transformation* S4
*Bacterial Conjugation* S4
*Plasmid Construction* S4
*Bioinformatic Analysis of the Pazamine Gene Cluster* S5
*Generating P. azotoformans Knockout Strains.* S5
*HILIC/QTOF-MS Analysis of Metabolites* S5
*Comparative Metabolomics* S6
*Stable Isotope Labeling* S6
*Amino Acid Fmoc-Derivatization and QQQ-MS/MS Analysis of Metabolites* S6
*Density functional theory (DFT) calculations* S7
*NMR spectroscopy* S7
*Isolation of Pazamide (**2**).* S7
*Pazamide Dimethyl Ester (**3**)* S8
*PazB Modeling and Substrate Docking* S8
*Germination of Arabidopsis thaliana* S8
*Plant Inoculation Assays* S8
*Protein Expression and Purification* S9
*In Vitro Assays for PazB Activity* S10


#### Supplementary Results


***Table S1.** Strains, plasmids, oligonucleotides, and DNA sequences used in this study* S11
***Figure S1.** SHMT and PazB sequence similarity network* S19
***Figure S2.** Serine hydroxymethyltransferase (SHMT) sequence analysis* S20
***Figure S3.** Gene neighborhoods surrounding the pazRABC cluster* S21
***Table S2.** Summary of PazB variants* S22
***Figure S4.** Metabolomic analysis with MS-DIAL* S24
***Figure S5.** 1D-NMR characterization of pazamide dimethyl ester (**3**)* S25
***Figure S6.** 2D-NMR characterization of pazamide dimethyl ester (**3**)* S26
***Figure S7.** 1D-NMR spectrum of pazamide (**2**)* S29
***Figure S8.** Confirmation of the PazA product* S30
***Figure S9.** Structural and sequence analysis of PazB active site* S31
***Figure S10.** Arginine/ornithine succinyltransferase can succinylate pazamine* S32
***Figure S11.** Arabidopsis seedling growth phenotype after bacterial inoculation* S33
***Figure S12.** Deuterium labeling supports PazB-mediated cyclization* S34



#### References S35

#### Author Contributions S37

### Materials and Methods

**Commercial Materials.** L-Lysine, iron(II) sulfate heptahydrate, ammonium formate (LC-MS grade), Dowex 50WX8 hydrogen form, irgasan (Irg) α-ketoglutarate (αKG) disodium salt dihydrate, lysozyme, 2-mercaptoethanol (βME), polyethyleneimine (PEI), pyridoxal 5′-phosphate (PLP), sodium dodecyl sulfate, sodium ascorbate, sucrose, trimethylchlorosilane (TMSCl) and were purchased from Sigma-Aldrich (St. Louis, MO). Acetonitrile, agarose, bromophenol blue, carbenicillin disodium salt, diethyl ether, dithiothreitol (DTT), deoxynucleotides (dNTPs), ethylenediaminetetraacetic acid disodium salt dihydrate (EDTA), ethyl acetate, formic acid, hydrochloric acid, gentamicin sulfate (Gm), HEPES, imidazole, kanamycin sulfate (Km), methanol, GeneRuler 1 kb Plus DNA Ladder, PageRuler Plus Prestained Protein Ladder, sodium chloride, sodium hydroxide, were purchased from Thermo Fisher Scientific (Waltham, MA). Acetic acid, dimethylsulfoxide, glycerol, LB Miller Agar, LB Miller Broth, nutrient broth, magnesium chloride hexahydrate, Terrific Broth, were purchased from EMD-Millipore (Burlington, MA). Absolute ethanol was from VWR International (Radnor, PA). InstantBlue Protein Stain was from Expedeon (San Diego, CA). Isopropyl ß-D-1-thiogalactopyranoside (IPTG) was from Santa Cruz Biotechnology (Dallas, TX). Bradford assay reagent concentrate and ethidium bromide were purchased from Bio-Rad Laboratories (Hercules, CA). Deuterium oxide, L-lysine dihydrochloride ($^{13}C_6$, 99%; $^{15}N_2$, 99%), L-lysine dihydrochloride (4,4′,5,5′-$d_4$), L-lysine dihydrochloride (2,3,3′,4,4′,5,5′,6,6′-$d_9$), and disodium succinate (2,2′,3,3′-$d_4$) were purchased from Cambridge Isotope Laboratories (Tewksbury, MA). Phusion polymerase, Phusion HF buffer, all restriction enzymes, restriction enzyme buffer (CutSmart), and Taq ligase were from New England Biolabs (Ipswitch, MA). T5 exonuclease was from Epicentre (Madison, WI). Ni-NTA agarose resin and DNA purification kits were purchased from Qiagen (Redwood City, CA). Oligonucleotides and gBlocks gene fragments were synthesized by Integrated DNA Technologies (Coralville, IA) or Twist Bioscience (South San Francisco, CA). All chemicals were used as purchased without further purification.

**Bacterial Strains and Culture.** *Escherichia coli* DH10B-T1$^R$ was used for cloning, *E. coli* BL21 Star (DE3) was used for protein expression and purification, and *E. coli* SM10 was used to transfer *oriT*-containing plasmids via bacterial conjugation. Unless otherwise stated, *E. coli* liquid cultures were grown at 37°C and 200 RPM on LB Miller broth with appropriate antibiotics, while single colonies were grown on LB Miller agar plates (1.5% *w/v*) at 37°C. *Pseudomonas azotoformans* (DSM 18862) was purchased from the German Collection of Microorganisms and Cell Cultures (DSMZ). *Pseudomonas azotoformans* strains were grown in LB Miller broth at 30°C and 200 RPM with appropriate antibiotics, while single colonies were grown on LB Miller agar plates (1.5% *w/v*) at 30°C. *P. azotoformans* strains were stored as glycerol stocks at -80°C and streaked to single colonies before cell culture for various experiments.

**Bacterial Transformation.** To 100 μL of chemically competent *E. coli*, plasmid DNA, 20 μL of KCM solution, and water up to a final volume of 200 μL were added. The mixture was incubated on ice for 30 min prior and then incubated at 42°C for 90 s. The cells were returned to ice for 2 min, 600 uL of LB Miller broth was added, and the cells were recovered at 37°C for 1 h before plating on selective media. *Pseudomonas azotoformans* was transformed via electroporation. A freezer stock of *P. azotoformans* was streaked out onto LB Miller agar and incubated overnight at 30°C. A single colony was used to inoculate 5 mL of LB Miller broth and the starter culture was grown at 30°C, 200 RPM overnight. The following morning, 25 mL of LB Miller broth was inoculated with 1% (v/v) starter culture and grown at 30°C, 200 RPM until $OD_{600}$ of 0.4 – 0.6. The culture was chilled in an ice bath for 15 min and then centrifuged at 5000 × *g* for 10 min. The supernatant was gently decanted, and the cell pellet was washed with 10 mL of chilled electroporation solution (1 mM HEPES, 1 mM $MgCl_2$, pH 7.2). The cells were once again pelleted at 5000 × *g* for 10 min and then resuspended in 1 mL of electroporation solution. The cells were washed as such two more times before resuspension in 500 μL of electroporation solution. Then, 100 μL of electrocompetent *P. azotoformans* was added to a chilled 2 mm gap width electroporation cuvette along with 50 - 100 ng of plasmid DNA. The cells were electroporated with a Bio-Rad MicroPulser on the Ec2 setting (2.5 kV, 5 ms). The cells were immediately resuspended in 800 μL of LB Miller broth and recovered at 30°C, 200 RPM for 90 min prior to plating on selective media and incubating at 30°C overnight.

**Bacterial Conjugation.** Plasmid DNA was also introduced to *P. azotoformans* by conjugation. First, *E. coli* SM10 was transformed with the plasmid to be conjugated by the heat shock method described above. Then, a freezer stock of *P. azotoformans* was streaked out onto LB Miller agar and incubated overnight at 30°C. A single colony of *E. coli* SM10 and *P. azotoformans* were separately used to inoculate 5 mL of LB Miller broth. When the cultures each reached an $OD_{600}$ of 0.8-1.0, they were mixed 1:1 to a final volume of 1 mL. The combined cells were then collected by centrifugation at 18,000 × *g* for 1 min and the supernatant was removed. The cell pellet was resuspended in 200 μL of LB Miller broth and plated directly in the center of an LB Miller agar plate and incubated overnight at 30°C. The next day, the bacterial mating mixture was scraped from the plate, resuspended in 1 mL of LB Miller. The resuspended mating mixture was diluted 1:10 and 1:100 in LB Miller broth and plated on LB Miller agar plates containing Gm (30 μg $mL^{-1}$) and Irg (20 μg $mL^{-1}$). After incubating overnight at 30°C, only transformed *P. azotoformans* remained.

**Plasmid Construction.** Gibson assembly and cloning by homologous recombination (HR) were routinely used to construct plasmids, using *E. coli* DH10B-$T1^R$ as the cloning host. Briefly, plasmids were first designed *in silico* via Benchling (Benchling, San Francisco, CA). The plasmid backbones were digested with the selected restriction enzymes and purified via the Qiagen PCR Cleanup Kit (Qiagen, Redwood City, CA) or amplified by Q5 polymerase (New England Biolabs, Ipswitch, MA). The insert amplicons were generated with either Phusion polymerase or GoTaq polymerase via the primers listed in *Table S1* and purified as above. For Gibson assembly, the inserts and backbone were mixed in 1:1, 1:3, and 1:5 ratios in 5 µL volumes, mixed with 15 µL of Gibson master mix [*1*], and incubated at 50°C for 1 h. The Gibson mix was transformed into chemically competent *E. coli* DH10B-T1$^R$ by heat shock, plated on the appropriate selective media, and incubated at 37°C overnight. For HR cloning, 100 ng of plasmid was mixed in a 1:2 molar ratio with insert and directly transformed into *E. coli* DH10B-T1$^R$ by heat shock, plated on the appropriate selective media, and incubated at 37°C overnight. Positive colonies were identified by colony PCR. Plasmids were sequenced at Genewiz to confirm the identity of the inserted sequence (Genewiz, South Plainfield, NJ)**.**

**Bioinformatic Analysis of the Pazamine Gene Cluster.** The genes in the *pazRABCD* cluster were searched against the SwissProt database using BLAST [*2*] to identify similar proteins which have been characterized in the literature. The *pazB* gene was further investigated with the EFI-EST [*3*] web tools by generating a sequence similarity network (SSN) of the SHMT Pfam (PF00464) and PazB (A0A1V2JN15). Only sequences from bacteria and archaea were included. The SSN was further manipulated within Cytoscape [*4*] to generate additional SSNs.

**Generating *P. azotoformans* Knockout Strains.** Genes were disrupted in *P. azotoformans* following the protocol from Hmelo et al [*5*]. Briefly, upstream and downstream genomic regions of the genes of interest were amplified from bacterial colonies of *P. azotoformans*. For every kilobase pair (kb) of DNA to be knocked out (*n*), the genomic homology arms of approximately 500 + *n**50 bp were amplified by PCR. Homology arms were designed to generate an in-frame deletion of the genomic region of interest, resulting in a small peptide product. Primers for these amplicons can be found in *Table S1*. These homology arms were cloned into the pEXG2 cloning vector by Gibson assembly as described above and were delivered to *P. azotoformans* by bacterial conjugation, as described above. A single colony of conjugated *P. azotoformans,* which has undergone a single crossover event, was picked and streaked onto a nutrient broth plate containing 1.5% (*w/v*) agar and 5% (*w/v*) sucrose and incubated at 30°C overnight. The following day, single colonies were screened by colony PCR for individuals which have successfully undergone a double crossover event and have lost the gene of interest.

**HILIC/QTOF-MS Analysis of Metabolites.** Samples were analyzed via hydrophilic interaction liquid chromatography (HILIC) coupled with high-resolution quadrupole time-of-flight mass spectrometry (QTOF-MS). Samples were analyzed using an Agilent 1290 UPLC (Santa Clara, CA) on a SeQuant ZIC-pHILIC (5 µm, 2.1 × 100 mm; EMD-Millipore, room temperature), with buffer A (90% acetonitrile, 10% water and 10 mM ammonium formate) and buffer B (90% water, 10% acetonitrile and 10 mM ammonium formate) comprising the mobile phase. The column was kept at 40°C. A linear gradient from 100% to 60% buffer A over 17 min, followed by a linear gradient from 60% to 40% buffer A over 8 min was performed at a flow rate of 0.2 mL min$^{-1}$. Mass spectra were acquired in positive-ionization mode using an Agilent 6530C QTOF mass spectrometer (gas temperature = 300°C, nebulizer = 35 psi, capillary voltage = 3500 V, nozzle voltage 500 = V, fragmentor voltage = 175 V, skimmer = 65 V, Oct 1 RF Vpp = 750 V).

**Comparative Metabolomics.** Starting from single colonies isolated from glycerol stocks, *Pseudomonas azotoformans* pPazAB and *P. azotoformans* pMMPc-Gm were grown in triplicate, as described above, in LB Miller broth (25 mL) containing Gm (30 μg $mL^{-1}$) in 250 mL baffled shake flasks. After 24 and 72 h, 2 mL of cells were removed from the culture and centrifuged at 21,300 × *g* for 1 min. The supernatant was separated from the cell pellet and mixed 1:1 with 1% (*v/v*) formic acid in methanol. The wet weight of the cell pellet was measured and resuspended in 1% formic acid in methanol (5 μL $\mu g^{-1}$ of wet cell pellet). The methanolic samples were vortexed to homogeneity and then centrifuged at 21,300 × *g* for 10 min. The supernatants were sampled without disturbing the pellet and analyzed by HILIC/QTOF-MS as described. The resulting data files (.d) were exported and converted to .abf files (Reifycs Analysis Base File Converter; https://www.reifycs.com/abfconverter/) and the resulting files were imported into and analyzed with MS-DIAL [*6*], a software package for untargeted metabolomics analysis For peak identification, an MS1 tolerance of 0.01 Da was used, along with a minimum peak height amplitude of 5000. For alignment, a retention time tolerance of 0.15 min and a MS1 tolerance of 0.01 Da was used. For chromatographic features to be included in the same group for analysis, >50% of the samples had to contain the chromatographic feature of interest (%N detected in at least one group). In the resulting alignment file generated by MS-DIAL, data in the ion table were restricted to *m/z* values between 100 and 300 and then sorted by fold-increase to identify differentially biosynthesized metabolites which were only present in the pPazAB overexpression condition. Metabolites with both high fold-increases and large signal-to-noise ratios (>20) were recorded as molecules of interest.

**Stable Isotope Labeling.** Starting from a single colony isolated from a glycerol stocks, *Pseudomonas azotoformans* pPazAB was grown in LB Broth (25 mL) containing Gm (30 μg $mL^{-1}$) in 250 mL baffled shake flasks as described above. After reaching $OD_{600}$ of 0.8-1.0, 5 mL of the culture was transferred to a 50 mL glass culture tube and fed 20 mg of isotopically labeled substrate ($^{13}C_6$,$^{15}N_2$-L-lysine, 4,4′,5,5′-$d_4$-L-lysine, 2,3,3′,4,4′,5,5′,6,6′-$d_9$-L-lysine, 2,2′,3,3′-$d_4$-succinate) to a final concentration of 4 mg $mL^{-1}$. After another 24 h of growth, the metabolome was extracted from the cell pellet as described and analyzed by the general HILIC/QTOF-MS method.

**Amino Acid Fmoc-Derivatization and QQQ-MS/MS Analysis of Metabolites.** Unlabelled and isotopically labelled metabolomic samples were grown as described above. The cellular metabolome was extracted as described above except using pure methanol, without formic acid. To 50 μL of the metabolomic extract, we added 12.5 μL of 200 mM, pH 8 sodium borate buffer and 10 μL of 10 mM Fmoc chloride in dichloromethane. The mixture was briefly vortexed and allowed to react at room temperature for 10 min. The sample was then analyzed via reverse phase chromatography coupled with triple-quadrupole tandem mass spectrometry (LC-QQQ-MS/MS). Samples were analyzed using an Agilent 1290 UPLC (Santa Clara, CA) equipped with an Agilent Poroshell 120 EC-C18 column (2.7 μm, 2.1 × 50 mm). Solvent A (0.1% formic acid in water) and solvent B (acetonitrile) comprised the mobile phases. Solvent A was kept at 100% for 1 min, followed by a linear gradient from 0% solvent B to 100% solvent B for 5 min. The mobile phase was then held at 100% solvent B for 1 min, before returning to 100% solvent A over the next min. The flow rate was 0.6 mL $min^{-1}$. Mass spectra were acquired in positive mode on an Agilent 6460 QQQ MS (300°C, gas flow = 5 L $min^{-1}$, nebulizer = 45 psi, capillary voltage = 3500 V, nozzle voltage = 500 V fragmentor = 135 V, the collision energy = 10, cell accelerator voltage = 7). The product ions (*m/z* = 152.1 $[M+H]^+$) were detected in MRM mode from precursor ions resulting from Fmoc-derivatized, perdeuterated compounds **1** and **7** (*m/z* = 374.2 $[M+H]^+$).

**Density functional theory (DFT) calculations.** All DFT calculations were performed in Gaussian 16, Revision B.01 [*7*]. Geometry optimizations were carried out using the hybrid three-parameter Becke´s (B3LYP) functional [*8*] with the empirical correction to dispersion (+GD3) and the def2-SVP basis set, with water as the solvent in the Self-Consistent Reaction Field (SCRF). Frequency calculations on the optimized geometries were performed in the same level of theory (B3LYP/GD3/def2-SVP/SCRF=water) to obtain the enthalpy and entropy corrections for the calculated energies.

**NMR spectroscopy.** Experiments were recorded on a Bruker Avance II spectrometer operating at 900 MHz and at 298 K, a Bruker Avance III spectrometer operating at 600 MHz and at 298 K, and a Bruker Avance I spectrometer operating at 700 MHz and at 298K. The 900 MHz instrument was equipped with a CP TXI cryoprobe, the 600 MHz instrument was equipped with a 5 mm 1H/BB Prodigy CryoProbe, and the 700 MHz instrument was equipped with a 5 mm triple resonance 1H/13C/15N TXI cryo-probe. All instruments were controlled using Topspin (Version 3.2) software. Data were processed using Topspin (Version 3.2) by zero-filling once in each dimension, followed by apodization, Fourier transformation, and phasing. Spectra were analyzed using Mnova software (Mestrelab Research, Escondido, CA, USA).

**Isolation of Pazamide (2).** A glycerol stock of *P. azotoformans* pPazAB was streaked onto an LB Gm plate and incubated at 30°C for 36 h. A single colony was used to inoculate 50 mL of LB Gm media and an overnight culture was grown at 30°C, 200 RPM. The overnight culture was used to inoculate 6 × 500 mL (3 L) of LB media in 2.5 L baffled shake flasks and grown for 24 h at 30°C, 200 RPM. The cell pellets were harvested by centrifuging at 8000 × *g* for 5 min and extracted with 2 × 50 mL of 80% (*v/v*) methanol in water with 1% (*v/v*) formic acid. The pellets were resuspended to homogeneity by vortexing. The extract was centrifuged at 8,000 × *g* for 15 min and the supernatant was collected and dried by rotary evaporation. The extract was purified by preparative HPLC using an Eclipse XDB-C18 column (5 µm, 9.4 mm × 250 mm, Agilent) on an Agilent 1260 HPLC with 100% water run isocratically as the mobile phase (5 mL $min^{-1}$) for 15 min. Fractions containing pazamide were identified by LC-MS and were pooled and dried to give a semi-pure extract. The semi-pure extract was resuspended in 50% ACN/50% $H_2O$ and further purified by preparative HPLC using a ZIC-HILIC column (5 µm, 21.1 mm × 50 mm, Supelco) on an Agilent 1200 HPLC with buffer A (10% ACN in water, 10 mM ammonium formate) and buffer B (90% ACN in water, 10 mM ammonium formate) comprising the mobile phase (5 mL $min^{-1}$). Buffer A was increased linearly from 10% to 40% at 8.08 min, 40% to 60% at 12.11 min, 60% to 70% at 18.17 min, and 70% to 10% at 20.19 min. Fractions containing 2 were identified by LC-MS, pooled, and evaporated to yield 22 mg of pazamide as an off-white powder. $^{1}$H NMR (700 MHz, $D_2O$) δ 3.09 (hept, J = 7.4 Hz, 2H), 2.62 – 2.57 (m, 2H), 2.51 (ddd, J = 8.2, 6.6, 2.0 Hz, 2H), 2.13 – 2.04 (m, 1H), 2.01 – 1.91 (m, 1H), 1.47 – 1.37 (m, 2H), 1.28 – 1.24 (m, 1H). $^{13}$C NMR (151 MHz, $D_2O$) δ = 178.07, 176.31, 176.26, 39.38, 38.35, 30.60, 30.10, 26.69, 24.39, 20.66. Found QTOF/MS *m/z* = 245.1130 $[M+H]^+$ (Calculated *m/z* for $C_{10}H_{17}N_2O_5$ = 245.1132, -0.81 ppm error).

**Pazamide Dimethyl Ester (3).** Pazamide **2** was methyl esterified [*9*] to facilitate structural elucidation. The product was stirred in acidic methanol prior to purification with HPLC using a ZIC-HILIC column (5 µm, 21.1 mm × 50 mm, Supelco) on an Agilent 1200 HPLC with buffer A (10% ACN in water, 10 mM ammonium formate) and buffer B (90% ACN in water, 10 mM ammonium formate) comprising the mobile phase. Buffer A was increased linearly from 10% to 40% at 8.08 min, 40% to 60% at 12.11 min, 60% to 70% at 18.17 min, and 70% to 10% at 20.19 min. Fractions containing 3 were identified by LC-MS, pooled, and evaporated to dryness via rotary evaporation and high vacuum. $^{1}$H NMR (900 MHz, $D_2O$) δ 3.72 (s, 3H), 3.68 (s, 3H), 3.06 (td, J = 7.1, 2.9 Hz, 2H), 2.66 (t, J = 6.7 Hz, 2H), 2.59 – 2.51 (m, 2H), 2.05 (dp, J = 12.8, 6.5 Hz, 1H), 1.95 (tt, J = 15.0, 7.2 Hz, 1H), 1.59 (dtd, J = 9.6, 8.4, 6.1 Hz, 1H), 1.54 (dd, J = 8.3, 5.3 Hz, 1H), 1.42 (dd, J = 9.6, 5.3 Hz, 1H). $^{13}$C NMR (226 MHz, $D_2O$) δ 179.14, 178.13, 175.60, 55.73, 54.94, 41.60, 39.61, 32.56, 31.60, 30.83, 26.82, 24.69.

**PazB Modeling and Substrate Docking.** The protein sequence of PazB (UniProt: A0A1V2JN15) was supplied to the AlphaFold 2 Colab server, ColabFold [*10*], to generate a structural prediction of a PazB homodimer using the default parameters. Substrate models were generated from SMILES strings at the CACTUS server (http://cactus.nci.nih.gov/translate/). Substrate and protein models were prepared for docking and used in docking experiments with AutoDock Vina using the default parameters [*11*, *12*]. Generated poses were compared to a substrate-bound *E. coli* GlyA (PDB: 1DFO) crystal structure [*13*] to identify catalytically relevant poses before further analysis.

**Germination of *Arabidopsis thaliana*.** *Arabidopsis thaliana* (L.) Heynh Columbia 0 (Col-0) ecotype seeds were sterilized with 20% (*v/v*) bleach for 10 min before washing five times with sterilized water. The seeds were resuspended in 0.1% (*w/v*) agarose to allow for precise placement on 1/3 Murashige and Skoog, 1% (*w/v*) plant tissue culture agar plates prior to germination. After drying, the plates were sealed with micropore tape, wrapped in foil, and cold stratified at 4°C for 48 h. The plates were then unwrapped and placed in a growth chamber with a photoperiod of 16 h light (intensity of 130 µmol $m^{-2}$ $s^{-1}$), 8 h darkness, and set to 22°C for germination. The relative humidity was maintained at 60%.

**Plant Inoculation Assays.** The same plates containing the germinated *A. thaliana* seeds were transferred to a growth chamber and grown for 5-7 d at 22°C with a photoperiod of 16 h of light (intensity 130 µmol $m^{-2}$ $s^{-1}$) and a dark period of 8 h. The relative humidity was maintained at

60%. The seedlings were then inoculated with overnight liquid cultures of *P. azotoformans* that were adjusted with LB Miller broth to an $OD_{600}$ of 0.02. In a biosafety chamber, the plates were unsealed and the diluted bacterial culture (100 µL) was added to each individual *A. thaliana* plate in a line approximately 1 cm below the initial seedling placement. A negative control condition was prepared with nothing added. Plates were briefly left to dry before resealing with micropore tape and returning to the growth chamber. Plates were imaged on the day of inoculation and at 2 d intervals until 14 d post inoculation (dpi) using an EPSON V600 photo scanner with a black background for contrast. Images were then imported to RhizoVision [*14*] where measurements were taken according to the same process and settings across the experiment. Data frames were analyzed in Jupyter notebook. One-way and two-way ANOVA analysis with Tukey´s post hoc test were used for testing differences in treatments. Statistical significance is denoted as * ($p$-value < 0.05) or ** ($p$-value < 0.01).

**Protein Expression and Purification.** *E. coli* BL21 Star (DE3) cells were transformed, as described above, with the appropriate protein-expression plasmid. A single colony was used to inoculate an overnight culture of LB Miller broth and grown at 37°C, 200 RPM overnight. The next d, TB supplemented with the appropriate antibiotic was inoculated with 1% of the overnight culture and grown at 37°C, 200 RPM until the culture reached an $OD_{600}$ of between 0.8 and 1.0. At this point, the cells were cooled on ice for 20 min before being induced with IPTG (100 µM). The proteins were then expressed overnight (12 -16 h) at 16 °C, 200 rpm. Cell pellets were collected by centrifugation at 8,000 × *g* for 5 min at 4 °C. The supernatant was decanted, and cells were resuspended in 5 mL $g^{-1}$ of cell paste in lysis buffer (50 mM HEPES, 300 mM NaCl, 10 mM imidazole, 10 mM βME, pH 7.5) supplemented with 1 tablet per liter of cell culture of EDTA-free protease inhibitor cocktail (Roche). The resulting cell suspension was then lysed via sonication, using a Qsonica Q700 sonicator (Amplitude = 50, 5 s on, 25 s off, 2.5 min total process time, ½” tip). The lysate was then centrifuged at 13,500 × *g* for 20 min at 4°C to separate the soluble and insoluble fractions. DNA was precipitated in the soluble fraction with 0.15% (wt/vol) polyethyleneimine at 4 °C for 30 min, with periodic inversion. The precipitated DNA was then removed by centrifugation at 13,500g for 20 min at 4°C. The soluble lysate was incubated with Ni-NTA (0.5 mL $g^{-1}$ of cell paste) for 45 min at 4°C, resuspended and loaded onto a column by gravity flow. The column was washed with wash buffer (50 mM HEPES, 300 mM NaCl, 20 mM imidazole, 10 mM βME, pH 7.5) for 15–20 column volumes. The column was then eluted with elution buffer (50 mM HEPES, 300 mM NaCl, 300 mM imidazole, 10 mM βME, pH 7.5). Fractions containing the target protein were pooled according to absorbance at 280 nm and concentrated using an Amicon Ultra spin concentrator (10-kDa MWCO; Millipore). Protein was then exchanged into storage buffer (50 mM HEPES, 100 mM sodium chloride, 10% (vol/vol) glycerol and 10 mM βME, pH 7.5) using PD-10 desalting columns. For PazB and PazB-like enzymes, 20 µM PLP was included in the lysis and wash buffers. Final protein concentration was estimated according to the absorbance at 280 nm. For the PazB and PazB-like enzymes, those which were insoluble were also co-expressed with the chaperones DnaK, DnaJ, and GrpE. In this case, the plasmid pKJE7 (Takara Bio Inc.) was co-transformed along with the PazB construct. The expression medium also contained 1 mg $mL^{-1}$ L-arabinose. The procedure was otherwise unchanged.

***In Vitro* Assays for PazB Activity.** All reactions (100 μL) contained L-lysine · HCl (2 mM), disodium αKG (6 mM), sodium ascorbate (2 mM), $Fe(SO_4)_2 \cdot 7H2O$ (100 μM), PLP (50 μM) and sodium chloride (> 20 mM, from protein storage buffer) in 50 mM HEPES buffer, pH 7.5. Reactions were initiated by addition of purified PazA (5 μM) and allowed to proceed for 45 min at room temperature before addition of PazB variants to a final concentration of 10 μM. Additionally, PazB activity was also assayed in the presence of tetrahydrofolic acid (225 μM), divalent metals ($Zn^{+2}$, $Mn^{+2}$, $Co^{+2}$ – 1 mM), or by starting 4-chlorolysine standard (2 mM). Upon addition of a PazB variant, the reaction was allowed to proceed overnight at room temperature. The reaction was then quenched with 5 volumes of methanol, 1% formic acid and centrifuged at 21,300 × *g* for 10 min at 4°C to remove the protein from the solution. The sample was then analyzed with the HILIC/QTOF-MS analysis described above.

### Supplementary Results

##### Table S1. Strains, plasmids, oligonucleotides, and DNA sequences used in this study.

#### A. Strains

| Strain | Description | Source |
|---|---|---|
| *E. coli* BL21(DE3) Star | *F-ompT hsdSB (rB-, mB-) gal dcm rne131* (DE3) | ThermoFisher |
| *P. azotoformans* DSM 18862 | Wild-type | DSMZ |
| *P. azotoformans* ΔAOST | Δ*aruFG* | This study |

#### B. Plasmids

| Plasmid | Description | Source |
|---|---|---|
| pMMPc-Gm | GmR, *lacI*, rep, mob, P*c*(*Delftia acidovorans* C17), oriRSF1010 | [*15*] |
| pPazAB | pMMPc-Gm, *pazAB* | This study |
| pJTL1 | pMMPc-Gm, *halBpazB* | This study |
| pJTL3 | pMMPc-Gm, *halDpazB* | This study |
| pEXG2 | GmR, OriT, ColE1, sacB | [*16*] |
| pEXG2-AOST | *aruFG* flanking regions (750 bp US/DS), GmR, OriT, ColE1, *sacB* | This study |

#### C. Oligonucleotides

| Oligonucleotides | 5´-Sequence-3´ | Notes |
|---|---|---|
| pazAB_F | tgtaatgcaggtagcgaacccgtgcgagggattgaaatgaactacgtacttg | Genomic amplification of *pazAB* |
| pazAB_R | ctaaaagcttggctgcaggtcgacgtcctgggttaatgacttagcgc | Genomic amplification of *pazAB* |
| pMMPc_F | atccgtcgacctgcagccaag | PCR linearization of vector |
| pMMPc_R | aattcacgggttcgctacctgc | PCR linearization of vector |
| Up_aruF_F | gcataaatgtaaagcaagcttctgcaggactcgcgaagacctggc | Genomic amplification of upstream homology arm |
| Down_aruF_R | ttcgactcccgagcagccaccagcatggtgtcactc | Genomic amplification of upstream homology arm |
| Up_aruG_F | gagtgacaccatgctggtggctgctcgggagtcgaa | Genomic amplification of upstream homology arm |
| Down_aruG_R | acccgtggaaattaattaaggtaccgctcgctcggtttgaacagc | Genomic amplification of upstream homology arm |

#### D. DNA sequences

| Genomic DNA | Sequence (ORF) |
|---|---|
| *pazAB* from *P. azotoformans* | CGAGGGATTGAAATGAACTACGTACTTGATGAGGCCCGCCTGCAACAACATCACCATAAT<br>AATTTTACCGAAGAGAAACTGTTTTCCTTGCGTCATGAGTTTTCTCGCGACGGCTTTATA<br>AAACTGCGTAATATCGTCGATGATGAGCTGCGCAACTTCATCACGGCAGAAGTTAACAGT<br>TTGATTGATCATCAACTTGAACGTCGCGACTTGCATCTGGCAACTACCGATAACACACCG<br>CGCTACATGAGCGTGGTGCGCAGCGAGTTCATTGCTGAAAACAGTCCGTTGATCAACGCG<br>CTGTCGAAGTCCGAAGTACTGCTCGGTACTTTGTCCCTGATTGCAGGCACGCCAATCGTG<br>GCATCCGTCTCCAAGGACGAGGAATTCCTGATCACCAAACAGGAACGCAAGGGGGATACC<br>CACGGCTGGCACTGGGGCGATTACAGTTTTGCCCTGATCTGGATTATCGAAACACCCCCG<br>ATTGCCAAGGGCGGGATGCTGCAATGTGTGCCTCACACCTCTTGGGACAAATCCAACCCG<br>CGGATCCACGAACTGCTGTGCAGTAATCCGATTGCGACCTACGGCTTCAAGACGGGAGAT<br>ATCTATTTCCTGCGCACCGACACCACCCTGCACCGCACCATTCCGCTGAATGAAGATGCC<br>ACGCGGATCATCCTCAATATGACGTGGGCCGCAGAAAAAGATCTTGCCCGAAACCTTCAC<br>GGCAATGACCGCTGGTGGGAAGACAAGCACGCCGAGGCCGCAAAAAGCCTGACGTAGGAC<br>ACGTCTGCCTGGTTGTCAGTTGAACCCATGGGATGCACGATCATGCACGATATCCAATGT<br>CTCCAACGCGCCAATGCCTTGCTTGCACAGGCGCACTCTCATGCAGCCATGGCTGACCTG<br>GTCCGGGCGGCCGTGGCTCGAAATGACCAGTGGCGCGGGCAACAGTGCATCAATCTCGTG<br>GCTGCCGAGTCTCCAACCAGTCCCTCTGTGCGCGCGTTGCTGTCCAGTGAAGTCGGCACG<br>CGCGCCTCAGGTGGGCACATTGGGCGTGACAACCGGTTTTTCTCTGGCATGAAGAATATC<br>GACGAACTGGAATCGCTGTGTGTGGAGCTGTTGAAAACCACCTTGCGCGTCCGGCATGCC<br>GATCATCGATTGCTGGGCGGTATGGCCGCTGTGCTGGCGGCTTACACGGCATTGGCCCGT<br>CCAGGCGACAATGTCATGACCGTTCCGGTGATCAGGGGCGGTGATACCAGCAACCGAACC<br>AATGGACCTCCCGGGGTACGCGGGCTCAAGGTCTGGGATATTCCCTTCGTACACAAGACT<br>GCGGATATCGATCTCAACCGCTTCAGCGAGGTTGCCCAGGCCTTGAAACCTGCCGTCATC<br>GGGCTGGGCATGACGTTGACCCTGTTCCCCCTGCCCGTGGCAGACATCAAGGGGATCGTC<br>TCGCCCTGGGGAGGCAAGGTCTATTTTGACGCCGCCCATCAACTGGGCTTGATCAGCGCC<br>GGGTTGTTCCAAGACCCACTGGCCGAAGGGGCGGACCTGATGACCGGCTCTTCCGGAAAG<br>ACCTTCAGTGGCCCTCAGGGCGGCGTGATGGTGTGGAACGATGACGCGTTGACGGCACCT<br>GTGCATGAGGCGATTTTCCCCACCCTGACCGGCAGCCATCAGATCAACCGGGTTGCGGCG<br>CTCGCGGTGGCGGCGACCGAGATGCTGGCGTATGGGCCGGCCTATATGAAGCAAGTGGTG<br>AGCAACGCACAGGCGTTGGCCCGGCACTTGGACCTGCGGGGCGTGACGGCCTTTTATCGA<br>GACCAGGGCTACACCCGAACCCATCAGATCGTGATCGATTCAAAACCCTTCGCCAGCGGC<br>CGCGAGGCGGTGCAGCGCCTGGAGGCGGCGAACATCATCGCCAATGAAATGCCGCTGCCT<br>TGGGACGCCAATCCCTACAAGGAAACGGGCATTCGATTGGGCACTGTAGAAGTCACCCGG<br>CTAGGGATGAAAGAGCGGGAGATGGCGTGGATAGCCGAACAGATTGCCAAGGTGCTGTTG<br>CATAGGGAGGACCCCATGGCGGTCGCCAACGGTGTGATCGATTTCATGAAGAACCACAGG<br>ACGGTCTACTTCTGCCATGAAAATGGGCTGCCGCGCTAAGTCATTAACCCAGGA |

| gBlocks (IDT) | Sequence (ORF) |
|---|---|
| **SiHalB-PazB** | ATGCAGGTAGCGAACCCGTGAATTGCGAGGGATTGAAATGACCACGAATACGCCTACAACCG<br>GTGTCCGTACGCTCGACCAGGACGCATTCGACCGTCATCTCAAGGAATTCTCCGAGAACGCC<br>CCCATCCTCGAGTACTCGCAAGCGTTCCGCCGGGAAGGTTACGTCAAGCTCACCGGTCTGGT<br>CTCGGCGGATCTGTTCGCCGAGGTCGTCAGCGAAGTGGGCGACCTGCTGGACCAGCACGCCC<br>AGCGTATCGACATCGAACTCAAAGAGACCGGGAATTCGCCGCGCAGGATGCACACGGTCAGC<br>GCGGCCGACATCGCACGCGATTCGCGCTTGATTAGCGCGATCTACGACTCCGCCAGCATACG<br>CGACACCCTGGGCCGGATCGCTCAAGCCGACGTGCGTCCATGCCCCTGGGAAGGTGAGAAAT<br>ACGTCATCATTCGCCAGGACCAGCCGGGCGACACCCATGGCTGGCACTGGGGGGACTTTAGC<br>TACACACTGATTTGGATCATCCAGGCGCCGGGCCCAGAGGTGGGCGGTATGCTGCAGTGCGT<br>GCCGCATACCAATTGGGACAAGCAAAACCCGCGCGTACACGAGTACCTTCAGAACCACCCCA<br>TCCGCACTTATGCCAACGCTACCGGCGATCTGTACTTCCTGCGTTCGGATACCACCCTGCAC<br>CGTACGATCCCGTTGAGCCGCCCGGCAACCCGCATCATCCTCAACACCTGTTGGGCCAGCAC<br>CGCCGATGTCGAACGGCCTACGACGCACGAAACCATGGACGCGATGTTCTCCAGTTGAGACA<br>CGTCTGCCTGGTTGTCAGTTGAACCCATGGGATGCACGATCATGCACGATATCCAATGTCTC<br>CAACGCGCCAATGCCTTGCTTGCACAGGCGCACTCTCATGCAGCCATGGCTGACCTGGTCCG<br>GGCGGCCGTGGCTCGAAATGACCAGTGGCGCGGGCAACAGTGCATCAATCTCGTGGCTGCCG<br>AGTCTCCAACCAGTCCCTCTGTGCGCGCGTTGCTGTCCAGTGAAGTCGGCACGCGCGCCTCA |

| | |
|---|---|
| | GGTGGGCACATTGGGCGTGACAACCGGTTTTTCTCTGGCATGAAGAATATCGACGAACTGGA<br>ATCGCTGTGTGTGGAGCTGTTGAAAACCACCTTGCGCGTCCGGCATGCCGATCATCGATTGC<br>TGGGCGGTATGGCCGCTGTGCTGGCGGCTTACACGGCATTGGCCCGTCCAGGCGACAATGTC<br>ATGACCGTTCCGGTGATCAGGGGCGGTGATACCAGCAACCGAACCAATGGACCTCCCGGGGT<br>ACGCGGGCTCAAGGTCTGGGATATTCCCTTCGTACACAAGACTGCGGATATCGATCTCAACC<br>GCTTCAGCGAGGTTGCCCAGGCCTTGAAACCTGCCGTCATCGGGCTGGGCATGACGTTGACC<br>CTGTTCCCCCTGCCCGTGGCAGACATCAAGGGGATCGTCTCGCCCTGGGGAGGCAAGGTCTA<br>TTTTGACGCCGCCCATCAACTGGGCTTGATCAGCGCCGGGTTGTTCCAAGACCCACTGGCCG<br>AAGGGGCGGACCTGATGACCGGCTCTTCCGGAAAGACCTTCAGTGGCCCTCAGGGCGGCGTG<br>ATGGTGTGGAACGATGACGCGTTGACGGCACCTGTGCATGAGGCGATTTTCCCCACCCTGAC<br>CGGCAGCCATCAGATCAACCGGGTTGCGGCGCTCGCGGTGGCGGCGACCGAGATGCTGGCGT<br>ATGGGCCGGCCTATATGAAGCAAGTGGTGAGCAACGCACAGGCGTTGGCCCGGCACTTGGAC<br>CTGCGGGGCGTGACGGCCTTTTATCGAGACCAGGGCTACACCCGAACCCATCAGATCGTGAT<br>CGATTCAAAACCCTTCGCCAGCGGCCGCGAGGCGGTGCAGCGCCTGGAGGCGGCGAACATCA<br>TCGCCAATGAAATGCCGCTGCCTTGGGACGCCAATCCCTACAAGGAAACGGGCATTCGATTG<br>GGCACTGTAGAAGTCACCCGGCTAGGGATGAAAGAGCGGGAGATGGCGTGGATAGCCGAACA<br>GATTGCCAAGGTGCTGTTGCATAGGGAGGACCCCATGGCGGTCGCCAACGGTGTGATCGATT<br>TCATGAAGAACCACAGGACGGTCTACTTCTGCCATGAAAATGGGCTGCCGCGCTAAGATCCG<br>TCGACCTGCAGCCAAGCTT |
| **PkHalD-PazB** | AATGCAGGTAGCGAACCCGTGAATTGCGAGGGATTGAAATGAGCGAACAAACGTCCAGCTTG<br>GTTATTGAAGTAATGGAGCAGCAACTGGCCAAGCATTTCCAGGCTATTTTGCAAGATGAAAA<br>CCGGATGAAGCAGATTCGTAATGAATTTCGCCGAGACGGTTATTTCAACTTCAAGAACTTTT<br>CGTTTCTACCGAAAAGGATCTTGGAAAATGTCCACGCCGAAGTCCATGCGTTATTGGATGAA<br>TATTCGGTGCGTCGCGATGTTACTGTTCCGTCCACAGGCAATACCTACCGCAAGATGTACAA<br>CGTCAACCAACCGGAGATCGCCGAAGGCGGGACATTCATCCCTGCGCTCTACCAATCCGAGT<br>CGCTGCGTAAGTTCCTGGGCAACATCGCTGGCGATGATCTGGCGTCTTGCTGGGAGCAGGAG<br>CAATACCTGGTCACCAAGCTGAGCCACCCGGGCGATACCCACGGCTGGCATTGGGGGGATTA<br>CCCGTACACGATGATCTGGATCATCGAGGCGCCCGAGGACCCGGCGATTGGCGGCGTGCTCC<br>AGTGCGTGCCACACAGCGAATGGGACAAGCAGAACCCGCAGATCTGGCAGTACATCCTCAAT<br>AACCCGATCAAGTCCTACCACCATCTCAAGGGTGATGTGTATTTCCTCAAGTCCGACACCAC<br>GTTGCACCACGTCGTGCCGATTCAGCAGGAAACCACTCGGATAATTCTCAACACATGCTGGG<br>CCAGCGCCCATGACCGGCGAACCGATGTCGCCCACGAAAGCATCGAAGTGATCTGGGACACC<br>AAGGCCCGGACCACCGAAGAAGCCTGAGACACGTCTGCCTGGTTGTCAGTTGAACCCATGGG<br>ATGCACGATCATGCACGATATCCAATGTCTCCAACGCGCCAATGCCTTGCTTGCACAGGCGC<br>ACTCTCATGCAGCCATGGCTGACCTGGTCCGGGCGGCCGTGGCTCGAAATGACCAGTGGCGC<br>GGGCAACAGTGCATCAATCTCGTGGCTGCCGAGTCTCCAACCAGTCCCTCTGTGCGCGCGTT<br>GCTGTCCAGTGAAGTCGGCACGCGCGCCTCAGGTGGGCACATTGGGCGTGACAACCGGTTTT<br>TCTCTGGCATGAAGAATATCGACGAACTGGAATCGCTGTGTGTGGAGCTGTTGAAAACCACC<br>TTGCGCGTCCGGCATGCCGATCATCGATTGCTGGGCGGTATGGCCGCTGTGCTGGCGGCTTA<br>CACGGCATTGGCCCGTCCAGGCGACAATGTCATGACCGTTCCGGTGATCAGGGGCGGTGATA<br>CCAGCAACCGAACCAATGGACCTCCCGGGGTACGCGGGCTCAAGGTCTGGGATATTCCCTTC<br>GTACACAAGACTGCGGATATCGATCTCAACCGCTTCAGCGAGGTTGCCCAGGCCTTGAAACC<br>TGCCGTCATCGGGCTGGGCATGACGTTGACCCTGTTCCCCCTGCCCGTGGCAGACATCAAGG<br>GGATCGTCTCGCCCTGGGGAGGCAAGGTCTATTTTGACGCCGCCCATCAACTGGGCTTGATC<br>AGCGCCGGGTTGTTCCAAGACCCACTGGCCGAAGGGGCGGACCTGATGACCGGCTCTTCCGG<br>AAAGACCTTCAGTGGCCCTCAGGGCGGCGTGATGGTGTGGAACGATGACGCGTTGACGGCAC<br>CTGTGCATGAGGCGATTTTCCCCACCCTGACCGGCAGCCATCAGATCAACCGGGTTGCGGCG<br>CTCGCGGTGGCGGCGACCGAGATGCTGGCGTATGGGCCGGCCTATATGAAGCAAGTGGTGAG<br>CAACGCACAGGCGTTGGCCCGGCACTTGGACCTGCGGGGCGTGACGGCCTTTTATCGAGACC<br>AGGGCTACACCCGAACCCATCAGATCGTGATCGATTCAAAACCCTTCGCCAGCGGCCGCGAG<br>GCGGTGCAGCGCCTGGAGGCGGCGAACATCATCGCCAATGAAATGCCGCTGCCTTGGGACGC<br>CAATCCCTACAAGGAAACGGGCATTCGATTGGGCACTGTAGAAGTCACCCGGCTAGGGATGA<br>AAGAGCGGGAGATGGCGTGGATAGCCGAACAGATTGCCAAGGTGCTGTTGCATAGGGAGGAC<br>CCCATGGCGGTCGCCAACGGTGTGATCGATTTCATGAAGAACCACAGGACGGTCTACTTCTG<br>CCATGAAAATGGGCTGCCGCGCTAAGATCCGTCGACCTGCAGCCAAGCTT |
| **WP_169378548.1** | ATGACAATTATGCATGATATCCGTTGCCTTCAGCACGCGAGCACCTTATTAAAACAAGCACA<br>ATCGCATGCTGCGATGGCAGATCTGGTTCGCGCCGCTGTTGTACGCAATGAACAATGGCGCG<br>GGCAGCAGTGCATTAACTTGGTCGCCGCTGAGAGTCCAACCAGTCCTAGTGTTCGCGCTTTA<br>TTGTCTTCAGAGGTCGGAACCCGTGCCTCTGGAGGACATATCGGCCGCGACAATCGTTTTTT<br>CTCTGGAATGAAGAACATTGATGAGTTAGAATCACTGTGCGTTGAACTTTTGAAAACTACGT<br>TCCGCGTGCAGCACGCCGATCATCGCCTTATGGGAGGGATGGCTGCCGTCCTTGCCGCCTAC<br>ACCGCTCTTACCCGTCCTGAAGACAACGTGATGACCGTCCCGGTCATTCGTGGGGGGGATAC<br>CAGTAATCGTACAAACGGTCCACCGGGTGTCCGCGGTCTTAAAGTGTGGGACGTTCCCTTCG |

| | |
|---|---|
| | TACATAAGACAAGTGATATTGATCTTACACGTTTTAGCGAGGTCGCCCAGACGTTAAAGCCC<br>GCCGTAATTGGATTGGGGATGACTCTGACATTGTTCCCTCTGCCTGTCAGTGATATCAAGGG<br>AATTGTTTCTCCATGGGGCGGGAAGGTATACTTTGATGCTGCCCATCAGTTAGGGTTGATCA<br>GTGCAGGCTTATTCCAAGATCCGTTGGCAGAAGGTGCTGATCTGATGACTGGGTCAAGTGGG<br>AAGACCTTTTCAGGGCCTCAAGGTGGTGTGATTGTTTGGAATGAGACGGCCCTGACTGCTCC<br>AGTTCATGAAGCAATTTTCCCGACGTTGACGGGTTCCCACCAAATCAATCGTGTCGCGGCAT<br>TGGCGGTCGCTGCTACTGAAATGCTGGAATACGGGCAAGCGTATATGAAGCAAGTCGTAAGT<br>AATGCACAAGCATTGGCCACCCATTTGGATTTACGCGGCGTAACCGCATTTTACCGTGAACA<br>AGGTTATACGCGTACCCACCAAATCGTGATTGATTCGAAGCCCTTTGCTTCGGGTCGCGAAG<br>CCGTCGAGCGTTTGGAGGCCGCGAACATTATTACAAATGAGATGCCTCTGCCGTGGGACACG<br>GATCCATATAAAGAGTCGGGCATCCGTCTGGGTACCGTAGAAGTAACTCGTCTTGGTATGAA<br>AGAACGTGAGATGGCATGGATTGCGGACCAGATTGTATCTGTACTTTTGCATAAGGAAGATC<br>CAATGGCGGTAGCAAATGGAGTAATCGATTTTATGAAAAACTACCGTACGGTGTATTATTGT<br>CATGAGAACGGCTTACCGCGCATGCAGTGA |
| **PazB_PROSS-1** | ATGCATGATATTCAGGCATTACAACGTGCAAATGCTTTGTTAGCCCAAGCCCATTCCCATGC<br>CGCGATGGCGGACTTAGTACGCGCAGCTGTGGCCCGTAATGACCAATGGCGCGGCCAGCAGT<br>GCATTAATCTGGTTGCTGCAGAGTCTCCAACGTCTCCCTCTGTTCGTGCCTTACTTAGTTCA<br>GAAGTAGGTACGCGCGCGTCAGGGGGCCATATTGGTCGTGATAACCGTTTCTTTTCAGGTAT<br>GAAGAACATCGACGAGCTTGAGTCTTTATGTGTGGAGCTGCTGAAAACGATCTTTCGTGTGC<br>GTCACGCGGATCATCGCTTGCTGGGGGGTATGGCAGCCGTATTAGCAGCTTACACTGCGCTT<br>GCGCGTCCAGGTGATAATATCATGACGGTGCCGGTTATTCGTGGCGGCGACACCAGTAACCG<br>TACAAATGGTCCTGCAGGCGTACGCGGACTGAAGGTATGGGACATCCCTTTCGACCATAAGA<br>CATTTGATATTGATCTGGATCGCTTCTCGGAAGTGGCTCAGGCGCTTAAACCCAAGGTCATT<br>GGGCTTGGCATGACTTTGACATTATTCCCACTGCCGGTTGCAGATATCAAAGAAATTGTAGC<br>AGAGTGGGGAGCCAAGGTGTACTTCGATGCTGCCCACCAACTGGGACTGATTGCAGCAGGTT<br>TGTTTCAAGATCCCTTAGCAGAGGGCGCTGACCTGATGACGGGCAGTTCAGGCAAGACGTTT<br>TCTGGTCCACAAGGTGGGGTTATGGTCTGGAATGACGATGCGTTGACTGCCCCAATTCACGA<br>AGCAATTTTTCCGACACTGACCGGGTCTCATCAAATTAACCGTGTCGCTGCCTTAGCAGTGG<br>CAGCCACTGAAATGTTAGCCTATGGGGAGGCATATATGAAACAGGTGGTTTCCAACGCACAG<br>GCTCTTGCTCGTCATCTGGATGAACGTGGCGTGACAGTTTTTTATCGTGATCAAGGATACAC<br>TCGCACACACCAAATCGTTATCGACAGTGCCAAATTCGGATCTGGACGTGAAGCTGTGCAGC<br>GCCTGGAGGCAGCCAACATCATCGCCAACGAAATGCCCCTTCCCTGGGATGCGAATCCCTAT<br>AAAGAAACTGGCATCCGTTTGGGAACTGTCGAGGTCACTCGCCTTGGCATGAAAGAACGTGA<br>AATGGCCTGGATTGCGGAGATGATTGCTCGCGTGCTGCTTCATCGTGAAGATCCTATGGCGG<br>TGGCAAATGACGTTATTGACTTTATGAAGAATCACCGCACGGTGTACTTCGCTCACGAGAAT<br>GGTCTTCCTCGTTAA |
| **PazB_PROSS-3** | ATGCACGACATTCAGGCCCTTCAGCGTGCGAACGCCCTGTTAGCACAGGCCCATTCTCATGC<br>CGCAATGGCCGATCTTGTCCGCGCGGCAGTTGCTCGCAATGATCAGTGGCGCGGGCAACAAT<br>GCATCAACCTTGTCGCAGCGGAGTCGCCAACCTCCCCCTCTGTGCGCGCCCTTCTGTCTTCT<br>GAAGTAGGGACGCGCGCCTCCGGAGGCCATATTGGGCGTGATAATCGCTTCTTCTCGGGTAT<br>GAAGAACATTGATGAACTTGAAAGCCTTTGCGTCGAACTTCTGAAGAAAATCTTCCGCGTTC<br>GCCATGCCGACCATCGCTTATTAGGAGGGATGATGGCCGTTCTGGCTGCATATACAGCCTTG<br>GCCCGCCCCGGTGACAATATTATGACCGTACCGGTGATTCGCGGGGGAGATACATCGAATCG<br>TACGAATGGCCCCGCAGGAGTACGCGGGCTGAAGGTCTACGACATCCCTTTCGATCATGAAA<br>CGATGAATATTGACCTGGATCGCTTTTCGGAAGTCGCTCAAGCGTTAAAGCCAAAAGTTATT<br>GGACTGGGGATGACGTTGACTCTGTTTCCCCTTCCGGTCAAAGAGATTAAGGAAATTGTCGC<br>AGAATGGGGGGCGAAGGTATACTTTGACGCTGCTCACCAACTTGGTCTGATCGCCGCAGGAC<br>TGTTCCAGGACCCTTTAGCCGAAGGTGCTGATTTGATGACTGGTTCATCCGGCAAGACATTC<br>TCTGGTCCCCAAGGCGGGGTCATGGTATGGAATGACGATGCACTGACCGCCCCCATTCATGA<br>AGCGATCTTCCCGACGCTTACGGGCTCGCACCAAATCAATCGCGTTGCAGCACTGGCCGTAG<br>CTGCCACTGAGATGTTGGCGTATGGCGAGGCCTATATGAAACAAGTCGTTTCTAATGCGCAG<br>GCCTTGGCGCGCCACTTAGATGAGCGTGGGGTCCCCGTCTTCTACCGCGACCAGGGATACAC<br>GCGTACTCACCAGATTGTTATTGACGCCGCTGATTTTGGAAGTGGACGTGAGGCTGTGCAAC<br>GCTTGGAGGCAGCCAATATCATTGCCAACGAGATGCCGCTTCCTTGGGACGCAAATCCGTAC<br>AAAGAAACCGGCATTCGCCTTGGGACTGTGGAAGTCACACGTCTGGGGATGAAAGAGCGCGA<br>AATGGCTTGGATTGCGGAGCTGATCGCTCGCGTTCTTCTTCATCGCGAGGACCCCAAGGCTG<br>TTGCTAACGATGTCATTGACTTTATGAAGAACCACCGTACTGTCTATTACGCGCATGAAAAT<br>GGCTTGCCGCGTTAA |
| **PazB_PROSS-4** | ATGCACGATATTCAAGCGTTGCAGCGTGCAAATGCGTTATTAGCACAGGCACACTCACATGC<br>CGCGATGGCAGACCTGGTTCGCGCGGCGGTAGCCCGTAACGACCAGTGGCGTGGTCAACAAT<br>GCATCAATCTTGTTGCAGCTGAGTCTCCGACATCCCCAAGTGTGCGTGCCTTGCTTTCTTCA<br>GAAGTTGGGACGCGTGCGAGTGGCGGCCACATTGGGCGCGATAATCGCTTCTTTTCAGGTAT<br>GAAAAACATCGATGAGTTGGAGTCACTTTGCGTTGAGTTACTGAAAAAGATTTTCCGTGTTC |

| | |
|---|---|
| | GTCACGCCGATCATCGCTTGTTGGGGGGAATGTTAGCCGTGTTGGCCGCTTACACTGCCCTT<br>GCACGCCCAGGGGATAATATCATGGTTGTTCCTGTAATCAACGGAGGGGATACGAGTAACCG<br>CAAGAACGGACCGGCAGGCGTTCGTGGGTTAAAAGTGTACGATATCCCATTCGACCATGAGA<br>CCATGAACATTGACTTAGACCGTTTCTCAGAGGTAGCGCAGGCATTAAAGCCCAAGGTCATT<br>GGATTGGGGATGACTTTACTGCTGTTTCCCTTACCGGTCAAAGAAATTAAGGAAATTGTTGC<br>CGAATGGGGGGCCAAAGTATTATTTGACGCAGCCCACCAGTTAGGGCTGATTGCGGCTGGAG<br>TATTCCAGGACCCATTGGCAGAGGGCGCGGATTTAATGACAGGAAGCACCGGTAAGACTTTC<br>TCCGGCCCGCAGGGTGGAATCATGGTGTGGAACGATGACGCATTAACAGCTCCAATTCATGA<br>GGCTATCTTTCCGACGCTGACAGGAAGCCACCAGATTAACCGCGTTGCCGCGTTAGCTGTGG<br>CTGCGACAGAAATGTTGGCCTATGGCGAAGCATATATGAAGCAAGTCGTATCCAATGCCCAA<br>GCTCTGGCTCGTGCTTTGGATGAACGTGGAGTTACGGTGTTCTACCGTGATCAGGGGTACAC<br>ACGTACTCACCAGGTAGTTATTGACGCCGCGAAATTCGGTAGCGGGCGTGAAGCGGTGCAGC<br>GCTTAGAGGCCGCAAACATCATTGCAAATGAGATGCCCCTTCCTTGGGACGCAAACCCGTAT<br>AAAGAAACTGGTATTCGTCTTGGCACACAGGAAGTAACTCGTCTGGGCATGAAGGAGCGTGA<br>GATGGCTTGGATTGCTGAGCTTATTGCGCGTGTGCTTCTTCACCGCGAAGACCCAAAGAAGG<br>TGGCTAACGACGTGATTGAATTTATGAAGAATCATCGTACGGTGTACTATGCACATGAGAAT<br>GGACTTCCACGTTAA |

| **Synthetic genes** | **Sequence** |
|---|---|
| ***Legionella anisa* SHMT (A0A2Z3KPG0)** | ATGTTCACCACAGACCGTACTGTCCACGATCCTGCCCTTTTACAACAGGCGCAACGGGACTT<br>GATGCACTGTGAATCCTACAAGAGTATGCAGGACCTGTTATTAGACCTGATCCAGAAGAACG<br>ACACCTGGCGCACCAAGAAGTGTATTAACCTTGTTGCTGCGGAGTCTCCTATGTCCCGCCTG<br>GCGCGGTCACTGCTGGCATGTGATCTCTCGATGCGTACGGCCGGTGGGCACATCGGGAAGAA<br>GAACCGCAATTTCATGGCAACACAGTACATAGATCAACTGGAATCGATGTGTCATGTTCTGC<br>TTACTAATGTTTTCGAGTGTCACTACTGCGAACACCGCTTGCTGGGTGGTACTCAAGCCTGC<br>CACGTCGTCTACAGCTCTCTTATCAAACCCAACGATACGTTAATCACAGTTATGCCGGAGCA<br>CGGCGGCGACTCATCCAACTGCTCCCAATCTATGCCGGGCCTGTTGGGATTGAACATCATCC<br>CGATGCCGTTCCTGCCGGACCAATTAACAGTTGACCTTAACCAGTTAGAATACCTGGTTTGT<br>CGTTACCGCCCACAACTTATCGCATTAGGCTTCAGTATTTGTCTGTTTGAACAGCCGTTAAA<br>AGAAATCTGCACAATCGCACAAAAGTACAAGGTCAGAGTGTTCTACGATGCTGCTCATGAGC<br>TTGGCCTGATTGCGGGCAAGTGTTTCGCAAATCCTTTCAAACAGGGTATCACCGTCATGTCT<br>GGAAGTACTGGAAAGACGTTTTCTGGTCCGCAAGGTGGGTTGCTGCTGTGGGACGATGATGA<br>GCTGATCCAGCCGATCGCCTCTACTGTATTCCCTAACTTCGTTGGTACTTACCAATTGAACC<br>GGGTGGCCGCGTTAACACTTACGACGTTAGAGCTTCAGCAATACGGAGAGCGTTATATGGCT<br>CAAGTTGTACACAACGCCAAGGCCCTTGCCATGCACCTTTACGAGTTAGGCATCCCGGTCTT<br>CGCCAAAGAGAAGAATTACACTCAAACCCACCAGGTATTAATCAATGCAAAGCAATACGGCG<br>GTGGCTTCGTCGCTGCCAACCAACTCGAAGCCTGCAATATCATTGGTAATCACGTCAATATA<br>CCGGGAGACAATAAACCTCTCCAAGGCTTACGTATAGCCACGACGGAGATCACCCGTCGCGG<br>CATGAAGGAGACACATATGCGTAAAATCGCTGAGTTCATCTATCGAGCATTGGCCACTCACG<br>AGCCTGCACACTACATTGCACACGAGGTATCATTGCTGTCAGAGGCCTTCCAAGAAATTTAC<br>TATTGTTAA |
| ***L. wadsworthii* SHMT (A0A378LV02)** | ATGTTTATAACGGACTGGGCAATTGACGACTCGGCGCTGCTTCACCGTTCTCGTCGAGAGTT<br>TATGCATTGTCAGTCCTACGCAGAGATGCAGGATATGCTGATGTACCTGATTAAGAAGAACG<br>GCTCCTGGAGAAAGCAGTGTATAAACCTGGTAGCAGCAGAAAGCCCAATGAGCCAGATCGTT<br>CGCAGCCTCTTAGCTGACGATTTATCAATGCGGACGGCCAGCGGGTACATCGGAAAGAAGAA<br>TCGCTACTTCATGGCTACACAATACATTGACCAGTTCGAGAGCTTGTGCCATGTACTGCTTA<br>CTGACTTATTCGAGTGCAATTACTGCGATCACAGACTGATGGGTGGAACTCAAGCCTGTCAA<br>GTCGTTTACTCGACTTTAACCCAACCGAACGACACTTTGATCACCGTGATGCCTATGCACGG<br>TGGGGACTCCAGCAATTGCCGTCAATCGATGCCCGGCCTGCTGAATCTTAACATTGTTCCTA<br>TGCCGTTCTTACGGGATAATCTGACAATCGATTTGCATCAATTGGAGTATCTGTTAGATCGT<br>TACAAGCCGAAGTTAATATCATTGGGTTTCTCAATATGCCTCTTGGAGCAGCCTATCCGTGC<br>GATTTCCGCACTGTGCCAGAAGTTCAAAGTGCCAGTTTTCTACGATGCCTCACACGAGTTGG<br>GGCTTATTGCTGGGAAATGTTTCGCAAACCCATTTATCCAGGGGACAACCATTGTTAGTGGT<br>TCTACTGGGAAGACATTCTCGGGGCCACAAGGCGGCTTACTGTTATGGAACGATGACTATTT<br>GGTACAGCCGATAACTAATACGGTATTCCCGAACTTTGTGGGTACGTATCAATTAAATCGTG<br>TTGCAGCGTTAACCTTAGCGTCGTTAGAGATACAACAGTACGGTGAAGAGTACATGGCGCAA<br>GGGGTTCGGAATGCTAAGACTCTGGCGATGCAATTACATACGTTAGGGATTTCTGTATTTGC<br>ACAGGAGAAGAACTATACACAAACCCATCAAATTTTGATCGACGCGACCAAATATGGTGGTG<br>GTTTCGCCGCGGCAAACCGCCTGGAAGCGTGCCATATCATTACAAACCACGTAAATCTCCCG<br>GGTGACAAGGAAGGCTTCCGCGGAATACGCATAGCAACGACAGAGATGACTCGGCGCGGTAT<br>GAAGGAAAAGCATATGCAAGAGATAGTTCATCTTGTATACCGTGCTTTAGAGACAAGTGAGC |

| | |
|---|---|
| | CGGCGTCATCAATCGCTAAGGGTGCTAGCATTCTTAGTCAAGAATTTCAGAAGATTCACTAC<br>TGTTGA |
| ***L. cincinnatiensis* SHMT (A0A378IQC9)** | ATGAGTCATAAGGTTAAACTGGCTGATCCGGCGGTGTTACAACGTGCTAAGCAGCAGCTTCT<br>TCGTTATCCCACTTTCGCGGATAAGAGACACCATTTTATCGAGACAATACACTGCAATCAGG<br>AATGGCGGATGCAGCGCTGCATTAATTTAGTTGCGGCGGAAGGGCCTTTGTCCCCCACGGCA<br>CGGAGCTTATTGTCCGGTGACCTTTGTATCCGTACAGCTGGCGGACATATTGGCGCTAAGCA<br>ACGGTATTTCGCGGCTACTCAGTGGATTGACGAAATGGAAGCGTATACCTATGAATCAATCA<br>AATCTCTCTTCCAATGTCAATTCTGCGACTTTCGTTTGATCGGTGGTACGCAGGCTTGTCAA<br>GTTGTATACTCTCTTCTTACCAAGCCTGGGGATACCGTGATTGCGGTAATGCCGGAGCAAGG<br>CGGTGACAGTTCGCATAGTGAGCAAAGTATGCCTGGTTTGCTCCAACTCCATGTAATACCCA<br>TGCCTTTCTTGGAAGACAACCTTAGTATCGACCTTAACCGCCTTGAGCAGTTAGTAGACCAA<br>TACCACCCCACACTTATCTCTCTGGGGCTCTCTGTTTCGCTCTTTGAATTACCTCTGCAGGC<br>AATCCAAACGATAGCTCACCAACATGGCCGTGTGTTTTACGACGCAGCACACGAGTTAGGAT<br>TGATCGCTGGCCGGTGCTTCACAAATCCGTTCCTTCAAAAGATCGATGTGGTCTCTGGTTCC<br>ACGGGAAAGACCTTCTCAGGTCCACAAGGCGGTTTGCTGTTGTGGAACGACGAGACCCTGAA<br>TACCCACTTTAGTAGTCTGATCTTCCCGGGATTCGTAGGCACGTACCAATTAAACCGAGTCG<br>CTGCTTTAGGTCTTACAGCCCTGGAGTTTCTGGAGCACGGCAATGCTTACATGTCTCAAGTT<br>ATAAAGAATGCGCAGGCACTGGCTCAATCCCTGGACGCCTATGGTATGACAGTTTGGGCAAA<br>AGAGAAGGGATTCACGCAATCCCACCAAGTCCTGCTGGATATGACGGCCTACGCGGGTGGCT<br>GGAGAGCCACGCGCCGACTGGAACACTGTGACATTATCGGGAATCCAGTCTTCGTACCGGGA<br>ACAGCCTCAACCCTGCTTACAGGACTGCGTTTGGCGACCACTGAGATGACCCGTCGCGGAAT<br>GAAAGAGCCCGAGATGAAGATAATCGCAAACCTTATAGCTCGCGCCCTTTGCACAGACGAGC<br>CAAGTTCACAAATTGCGCACGACAGCAACTCTCTGGCTGCCCTTTTCCAAACCATTCACTAC<br>TGTTAA |
| ***E. coli* GlyA (P0A825)**<br>Y55A<br>A56S<br>E57G<br>Y65F<br>Y66F<br>S99M<br>H125D<br>L126T<br>T127S | ATGCTTAAACGCGAGATGAATATTGCCGATTATGATGCGGAACTGTGGCAAGCGATGGAGCA<br>GGAGAAAGTTCGACAAGAAGAACATATTGAATTGATCGCTAGTGAAAACTATACATCACCCC<br>GGGTGATGCAAGCACAAGGTTCACAGCTGACGAATAAAGCTTCTGGTGGATATCCAGGGAAA<br>AGATTTTTTGGCGGTTGTGAGTACGTAGACATCGTCGAACAGTTAGCTATCGATCGGGCAAA<br>AGAACTGTTCGGCGCGGATTACGCCAATGTGCAACCGCACAGTGGAATGCAGGCAAATTTTG<br>CTGTGTATACAGCTCTGTTGGAACCCGGGGATACAGTACTGGGGATGAATCTGGCACATGGC<br>GGGGATACCAGTCACGGCTCTCCAGTTAATTTTAGCGGGAAACTGTATAATATCGTTCCCTA<br>CGGCATAGACGCAACTGGGCATATAGATTACGCAGATTTGGAAAAACAGGCAAAAGAGCATA<br>AACCGAAAATGATCATCGGTGGTTTCTCAGCTTATTCAGGGGTGGTCGACTGGGCCAAAATG<br>CGTGAAATAGCCGATAGTATAGGTGCATATCTGTTTGTTGATATGGCACACGTCGCCGGTCT<br>CGTAGCCGCCGGCGTGTATCCCAATCCCGTTCCCCACGCTCACGTTGTGACAACTACCACAC<br>ACAAAACACTTGCAGGCCCTAGAGGAGGTCTCATTCTCGCGAAGGGCGGTTCTGAGGAATTA<br>TATAAAAAACTGAATAGCGCCGTATTTCCTGGCGGGCAGGGAGGCCCCCTGATGCATGTGAT<br>AGCAGGCAAGGCCGTTGCGCTTAAGGAAGCCATGGAACCAGAGTTCAAAACTTATCAGCAAC<br>AGGTAGCTAAGAATGCCAAAGCCATGGTCGAAGTTTTTTTGGAACGTGGTTATAAGGTTGTT<br>AGCGGAGGCACAGACAATCACTTATTCCTGGTTGATCTGGTGGATAAAAATCTGACGGGGAA<br>AGAGGCGGATGCTGCTCTTGGACGTGCAAACATAACCGTCAATAAAAATTCTGTCCCCAATG<br>ATCCGAAGTCCCCGTTTGTGACATCCGGCATACGTGTAGGTACCCCGGCAATTACACGCAGA<br>GGATTCAAAGAAGCTGAAGCCAAAGAATTGGCGGGTTGGATGTGTGATGTGCTGGACTCGAT<br>CAACGATGAAGCTGTGATCGAACGCATTAAAGGTAAAGTACTTGATATCTGCGCACGTTACC<br>CAGTTTATGCATAA |
| ***Salicibacter cibi* (JGI IMG: 2994814746)**<br>E79G<br>W88F<br>H150D<br>S151T<br>E79G | ATGTCAATGGTAGAAACGAATTACGAAGTTCATGATACCAAGGTCCTCAATTGGGCCAAAGA<br>GATTATTGAGGGTTCTCCATCTGAACAGTTGGTGCAACAGGAAGTGATTAATGCGGTGGAGC<br>GCAATGCTATTTGGCGGGGTCAGGAATGCTTGAATCTGCTGGCACCGGAAGCACCTACCAGC<br>CCTGCAGTCCGTCGCCTGCTGTCCGCAGAGGTAGGGACGCGGGCCGCGGGTGGACACATAGG<br>GTCACAGCAGCGTTTCTTTGCGGGGACGAAGTATATTGATGAAATAGAATCCTTATGCATTG<br>AGCTGCTGAAAAAGGTGTTCGATAGTAACTATGCCGATCATCGCTTGGTTGCTTCCATGATT<br>GGTAATATGGCAGTTTATACGGCTTTAACGGAGCCGGGGGATGATATCATGACCCTTAGCCA<br>GCCATTTGGCGGGGATTCAACTAACCGTAATGATGGCCCAGCGGGCGTACGAGGACTGAAAG<br>TGACCGATGTTCCTATGGATCCCGTTGAATTAGAGGTCGATCTGGATAAGTTCGCGGCTAAA<br>GCTCGGGAAAAGAAACCTAAAGTTGTTTCTCTGGGCGCCAGCATGACCTTGTTCCCATTTCC<br>TCTTAAGCAGATGAAGGAAATTGTGAGTGAATGGGAAGGGAAGATATTTTTTGACGGTGCCC<br>ACCAGCTGGGACTTATAGCTGGTGGCAAATTTCAGGATCCTTTGAATGAAGGCGCAGATGTG<br>ATGACGGGGTCCGCAGGTAAAACATTTAGCGGCCCTCAGTCCGGCATTATGGTATGGGACGA<br>TGTGAACTTAACTAAGCCGTTCACAGATACAATTTTCCCGACACTCGCGGCTACTCATCAAG<br>TAAATCGCGTGGCAGCGTTAGCCGTGAGCTGTGCAGAGTTTCTGGCGTTTGGAAAAGATTAT<br>ATGGAACAAATTGTAACGAACGCAAAAGCATTGGGTAATGCGCTGCATCAACGTGGAATAAG<br>TGTCCTGGGTTCAGATAAAGATTTCACAGAAACGCACCAGGTTATTCTTGACGTTAAACGCT<br>TTGGAGGAGGTTACGAGGTTGCTAATAAACTGGCGGAGGCTAATATTATTACCAATAAAAAC |

| | |
|---|---|
| | CTGATACCCGGTGATCGGCCTGAAGATTGGGATTATCCCAACGGACTGAGAATAGGCACGAC<br>AGAAATCACACGGAAGGGTATGAAGGAACCTGAGATGGAAATGATTGCCGACTATATCGATC<br>AACTGCTGAACAATCGGGAAACCCCCGAAAACGTGAAACTCAAAGTGCTGGACATGACGACT<br>TCATTCCAGAAAATTCACTATTGCTTT |
| **MPNN-PazB m1_21** | ATGCACGATCAGCAGTGCCTTGACAGAGCCAATGCTTTGCTGGCAGCCGCGCCGAGCCACGC<br>GGCTATGGCAGATTTAGTCCGTAGTGCCGTAGCTAGAAATGACCAATGGCGAGGCCAACAAT<br>GCATAAACCTCGTTGCAGCGGAGTCACCGACGAGTCCGAGCGTACGTGCACTGCTTTCAAGC<br>GAGGTAGGGACCCGTGCGAGTGGTGGTCATATTGGTCGCGACAACCGTTTCTTCTCGGGCAC<br>TAAGAACATCGACGAACTCGAGTCATTGTGTGTGGAGTTACTTAAAACACTGTTCCGTGTCA<br>AGCACGCAGACCACCGGCTGTTGGGCGGCATGGCCGCAGTTCTGGCCGCATACACCGCGCTC<br>ACTAAGCCAGGTGACAAGGTAATGTCAATCCCCGTGCGCTACGGCGGGGACACTTCAAATCG<br>TACGAATGGGCCGCCAGGGGTTCGCGGCCTGGAAGTACACGACATACCATTCAAGCCAGAAA<br>CGGGAGATATAGACCTTGACAAGTTCAAAGAGATAGCCGAGAAGTTGAAGCCCGCGGTGATC<br>GGTCTTGGTGCGACTCTGTTATTATTCCCTCTGCCTGTGGCGGAGATTAAGGACATAGTTAA<br>GCCCTGGGGTGGTAAGGTCCTTTACGACGCTGCGCACGCAGGCGGTTTGATAGCCGCAGGCC<br>TTTTCCAAGACCCACTGGCCGAAGGAGCTGACCTTATGACGGGCAGCTCGGGGAAAACCTTG<br>AGTGGGCCGCAGGGCGGGGTTATGGTCTGGAATGACGAAGCTTTAACTGTGCCGGTCCACGA<br>GGCGGTCTTCCCAACTTTGACAGGGTCCCACCAAATTAATCGCGTAGCCGCTATGGCGGTCG<br>CAGCCACCGAGATGCTGGCATACGGTCCTGCTTATATGACTCAAGTAGTTGCTAACGCCCGC<br>GCTCTGGCCAGAGCGCTGGACGAACGTGGAGTTACGGCATTCTACAAGGAGCGGGGCTATAC<br>GCAGACGCACCAGATTGTAGTTGACGCTAGCCCATTTGGTACAGGACGTGAGGCTGTTCAAG<br>CCTTAGAAGCCGCTAACATAATCGCAAACGAGATGCCATTACCATGGGACGAAGACATTGAT<br>AACCCATCTGGCATTCGATTAGGTACCGTCGAGGTGACAGCACTTGGTATGCGGGAAGATGA<br>CATGGGCGTTATCGCTGGCTGGATTGCGGACGTACTGTTAGACAAGAAGGACCCGAAAGAGA<br>TTGCCAAGAAGGTTATCGAGTTTATGAAGAAGCATCAAACTGTTTACTATAACCACGAGAAT<br>GGTCTGCCGCCTAAGCTTGCGGCCGCACTCGAGCACCACCACCACCACCACTGA |
| **MPNN-PazB m1_23** | ATGCACGATCAACAATGTCTGGACCGCGCCAACGCATTATTGGCGGCGGCACCATCTCACGC<br>AGCCATGGCTGATTTGGTTCGCAGTGCCGTTGCGCGTAATGACCAGTGGCGTGGACAACAAT<br>GCATCAACCTGGTCGCAGCGGAGTCACCCACGTCGCCGAGTGTGCGCGCTTTACTGTCGTCT<br>GAGGTAGGCACTAGAGCTTCAGGTGGCCACATCGGACGAGACAACCGCTTCTTTAGTGGGAC<br>GAAGAACATTGACGAGTTGGAAAGCTTATGCGTGGAACTGTTAAAGACGTTGTTTAAGGTAA<br>AACACGCTGACCACCGACTTCTTGGCGGCATGGCTGCGGTTCTGGCCGCATATACGGCCTTA<br>ACCTCGCCGGGCGACAATGTGATGAGCATTCCGGTTCGTTACGGCGGCGACACTTCCAACCG<br>CACCAACGGACCACCCGGAGTTCGTGGCCTTAAGGTGCATGACATCCCTTTCAAGCCTGAGA<br>CAGGAGATATTGACCTTGAACGCTTTGAAGAAATTGCCAAGAAGCTGAAGCCTGCGGTAATC<br>GGTCTGGGCGCTACGCTGCTGCTTTTCCCCTTACCAGTAGCTGAAATTAAGGATATAGTTAA<br>GCCATGGGGCGGGAAGGTCTTATTCGACGCTGCTCACGCGGGCGGACTTATCGCCGCTGGAT<br>TGTTTCAAGACCCCCTTGCAGAGGGCGCGGATCTCATGACTGGTAGCAGTGGAAAGACTCTG<br>TCCGGTCCTCAGGGCGGCGTGATGGTCTGGAATGATGAGGCGCTGACCACACCTGTGCATGA<br>GGCAGTCTTTCCCACACTGACCGGCTCCCATCAGATCAATCGGGTAGCAGCCTTGGCTGTTG<br>CCGCGACTGAGATGCTGGCATACGGGCCGGCGTACATGACCCAGGTGGTAGCTAATGCGCGC<br>GCGTTAGCTGCTGCGCTTGATGAGCGCGGCGTAACAGCATTTTATAAGGAACGGGGCTACAC<br>ACAAACGCATCAAATCGTTGTTGACGCGAGCCCCTTCGGGACCGGGCGGGAAGCTGTGCAAG<br>CACTCGAGGCAGCCAACATTATAGCTAATGAAATACCTCTGCCGTGGGACGAGGACATTGAA<br>AACCCGAGCGGCATTCGGTTGGGGACGGTTGAGGTTACGGCGTTAGGTCTCAGAGAAGAGGA<br>TATGGGTACAATTGCCGGATGGATAGCTGATGTTCTGCTGGATCGGCGTGACCCGGCGGAGA<br>TTGCTGCCGAGGTGGAAGAGTTCATGAAGGACCACCAAACTGTTTACTACAACCACGAGAAC<br>GGGTTGCCGCCGAAGCTTGCGGCCGCACTCGAGCACCACCACCACCACCACTGA |
| **MPNN-PazB m2_46** | ATGCATGATCAACAATGTCTTGACCGGGCTAACGCGTTGTTGGCAGCGGCTAGTAGTCACGC<br>CGCGATGGCCGATCTGGTACGGAGCGCTGTAGCCCGCAATGACCAATGGCGTGGACAAAAGT<br>GCGTCAATCTTGTGGCGGCTGAATCCCCGACATCGCCCTCTGTCCGGGCGCTTCTGAGCAGC<br>GAAGTTGGAACTCGCGCTTCCGGCGGCCACATTGGGCGCGATAATCGTTTCTTCTCAGGCAC<br>TAAGAACATTGACGAGCTTGAATCGTTATGCGTTGAATTATTGAAGACTACCTTAAACGTTG<br>CCTACGCCGATCACCGGCTGTTAGGTGGAATGGCTGCTGTCTTAGCCGCGTACACAGCATTG<br>GCCAGTCCGGGCGACGTCGTAATGACTATACCCGTTAAGTATGGTGGAGACACTTCGAACCG<br>TGAGAACGGCCCGCCTGGCGTCCGAGGACTCCAGGTCCATGATATACCTTTCGTACCTGATA<br>CTGGTGACATCGACTTAAACAAATTCGAAGAAGTGGCAAAGAAATTGAAGCCTGCGGTAATC<br>GGGCTTGGAAGAACCCTGTCACTGTTCCCGTTACCCGTCGCAGATATCAAGGGTATCGTGTC<br>CCCTTGGGGCGGGCAAGTATTCTTTGATGCCGCGCACCAACTGGGTCTGATCAGTGCAGGCC<br>TCTTCCAAGACCCACTGGCCGAGGGAGCGGATCTGATGACAGGTAGCAGTGGGAAAACATTT<br>GCAGGCCCGCAAGGTGGGGTTATGGTGTGGAACGACCCTGCTTTAACCGTCCCTGTCCACGA<br>GGCGATTTTCCCGACCCTTACCGGTTCTCATCAGATTAACCGTGTTGCCGCCTTAGCCGTGG<br>CAGCCACTGAAATGCTTGCGTACGGCCCGGCGTATATGACACAAGTAGTTCGGAATGCACAA |

| | |
|---|---|
| | GCCTTAGCCAGACACCTCGACGCCCGTGGTATCCGTGCGTTCTATCGAGAACGCGGGTATAC<br>GCAAACACATCAAATAGTTGTAGACGCTACGCCATTTGCCTCTGGGCGTGAGGCTGTTCAGG<br>CGCTCGAGGAAGCGAATATCATAGCGAACGAAATGCCGTTGCCATATGACGCCGACATCGAC<br>AAGGAGACAGGAATCCGGCTGGGCACGGTTGAGGTCACACGCCTTGGTATGCGGGAAGAGGA<br>GATGGCGTGGATTGCTGAGCAAATAGCCGCCGTACTCCTGGATAAGAAGGACCCACGTGAAG<br>TAGCTAAGAAGGTAGAGGAGTTTATGAAAGACCATCAAACAGTTTACTTTAACCATGAGAAC<br>GGGTTGCCGCCAAAGCTTGCGGCCGCACTCGAGCACCACCACCACCACCACTGA |
| **MPNN-PazB m2_48** | ATGCACGACCAGCAATGTCTTGACCGAGCGAATGCCTTATTGGCGGCCGCCCCGTCACATGC<br>GGCTATGGCAGACCTGGTTAGAAGCGCTGTGGCTCGCAACGACCAGTGGCGGGGTCAAAAGT<br>GTATAAATTTAGTTGCCGCCGAATCTCCAACTAGTCCGTCCGTCCGTGCTTTGCTTTCATCG<br>GAAGTAGGTACGCGGGCGAGTGGTGGGCACATAGGGCGGGACAATCGTTTCTTTTCAGGGAC<br>GAAGAATATTGATGAGCTGGAGTCTCTGTGTGTCGAGTTGTTAAAGACTACGCTGAACGTCA<br>AGTTCGCGGACCACAGATTGTTAGGTGGCATGGCTGCCGTGTTGGCGGCATACACCGCATTA<br>GCCGACCCTGGGGATGCGGTGATGACGGTACCAGTCCTTCAAGGTGGTGACACGTCCAACCG<br>TGCGAACGGACCACCGGGCGTTCGGGGTCTCCAAGTGCACGACATACCTTTCGTACCTGATA<br>CCGGTGACATCGATCTGAACGAGTTCCGCCGCGTGGCAGAGCGCTTAAAGCCTGCCGTGATC<br>GGGCTGGGGTTAACCTTGTCACTGTTCCCTTTACCCGTGGCGGACATTAAGGCAATCGTGTC<br>ACCTTGGGGCGGCCAGGTTTTCTTCGACGCCGCCCATCAGCTCGGGTTAATCTCGGCAGGAC<br>TTTTCCAAGACCCACTGGCGGAAGGTGCTGACCTGATGACTGGCTCTTCGGGGAAGACGTTT<br>GCGGGTCCCCAAGGCGGCGTGATGGTCTGGAACGACGAAGCCCTGACAACTCCGGTCCACGA<br>GGCGATTTTCCCTACCCTGACAGGCTCGCATCAGATCAATCGAGTGGCAGCCTTAGCTGTGG<br>CGGCGACCGAGATGCTGGCTTACGGTCCGGCATATATGACACAAGTTGTGCGAAACGCACAA<br>GCGCTTGCTCGTCATCTCCAGGAGCGCGGGATTCGTGCGTACTATGCAGAGCGTGGCTACAC<br>GCAGACACATCAAATCGTTATTGACGCATCTCCGTTCGCTACGGGACGTGAAGCTGTACAGG<br>CACTTGAAAAGGCCAACATTATCGCGAATGAAATGGCTCTGCCAACTGACGAAGACATATAC<br>CAGGAGACCGGTATTCGTTTAGGCACCGTCGAGGTCACTCGTTTGGGTATGCGTGAAGAGGA<br>GATGGCGTGGATTGCTGAGCAAATCGCAAAAGTTTTGCTTGACCGTGAGGACCCGAAACTTG<br>TCGCCAAGAAAGTGGAAGAGTTCATGAAGGATCACCAAACCGTCTACTTTAACCACGAGAAC<br>GGCTTGCCACCTAAGCTTGCGGCCGCACTCGAGCACCACCACCACCACCACTGA |

#### E. Gene Accession IDs

| Organism | Uniprot Accession | Gene | Function |
|---|---|---|---|
| *Pseudomonas azotoformans* | A0A1V2JQ47 | *pazR* | Transcriptional regulator |
| *Pseudomonas azotoformans* | A0A1V2JBT9 | *pazA* | L-Lysine 4-chlorinase |
| *Pseudomonas azotoformans* | A0A1V2JN15 | *pazB* | Carbocyclase |
| *Pseudomonas azotoformans* | A0A1V2JMT9 | *pazC* | Transporter |
| *Pseudomonas azotoformans* | A0A1V2JC74 | *pazD* | Transporter |
| *Escherichia coli* | P0A825 | *glyA* | Serine hydroxymethyltransferase |
| *Legionella anisa* | A0A2Z3KPG0 | *pazB* | Carbocyclase |
| *Legionella wadsworthii* | A0A378LV02 | *pazB* | Carbocyclase |
| *Legionella cincinnatiensis* | A0A378IQC9 | *pazB* | Carbocyclase |
| *Salicibacter cibi* | n/a | *pazB* | Carbocyclase (JGI IMG: 2994814746) |
| *Streptomyces iranensis* | A0A061AEA2 | *halB* | L-Lysine 5-chlorinase |

**Figure S1. SHMT and PazB sequence similarity network.** A sequence similarity network of serine hydroxymethyltransferases (PF00464) was merged with PazB (A0A1V2JN15) BLAST hits. Nodes represent protein sequences with greater than 80% sequence identity and are connected by edges at an alignment score threshold of 111, corresponding to a sequence identity of ~40%. Hits gathered from the SHMT Pfam are shown in blue, PazB BLAST hits are shown in magenta, and sequences which originate from both sources are in yellow. Bacterial sequences are shown as triangles while archaeal sequences are shown as circles. The sequence cluster which contains PazB-like enzyme nodes (SHMTs colocalized with a lysine halogenase) is expanded and PazB-like nodes are colored green. PazB-like enzymes are less than 40% identical to canonical bacterial SHMTs. Additionally, most of the sequences retrieved from the PazB BLAST search originate from archaea.

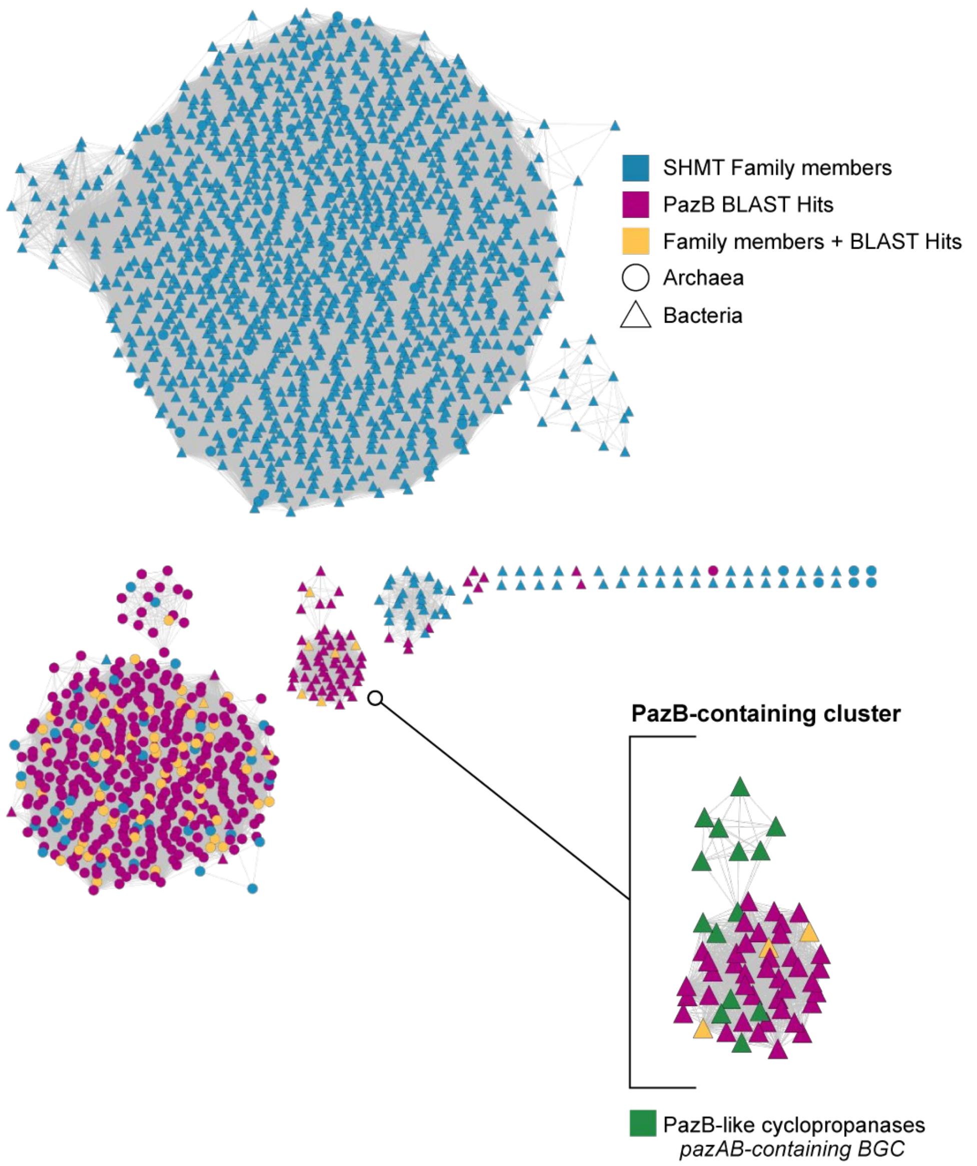

**Figure S2. Serine hydroxymethyltransferase (SHMT) sequence analysis.** (A) SHMT catalyzes the reversible conversion of 5,10-methylene-tetrahydrofolate (5,10-mTHF), glycine, and water to serine and THF. (B) A percent identity matrix of SHMTs shows that even between eukaryotes and prokaryotes, canonical SHMTs share reasonable percent identity (>40%). PazB from *P. azotoformans* and an SHMT-like enzyme from *L. anisa* share very little sequence identity with canonical SHMTs from bacteria and eukaryotes. The enzymes that are colocalized in biosynthetic gene clusters with amino acid halogenases do not contain highly conserved residues involved with folate binding and canonical SHMT catalysis. Using the numbering from *E. coli* GlyA, we can see that both the conserved E57 is glycine and the conserved N347 is valine or methionine in the PazB-like enzymes. Mechanistic and structural studies have shown E57 forms a necessary interaction with folate to catalyze the folate-dependent SHMT reactions [*17*] while N347 forms two hydrogen bonds with the pteridine ring of folate [*13*]. The lack of these residues suggests the PazB-like enzymes do not perform canonical folate-dependent chemistry. Additionally, the highly conserved H126 which π-stacks with the PLP ring [*18*] is instead an aspartate residue in the PazB-like enzymes.

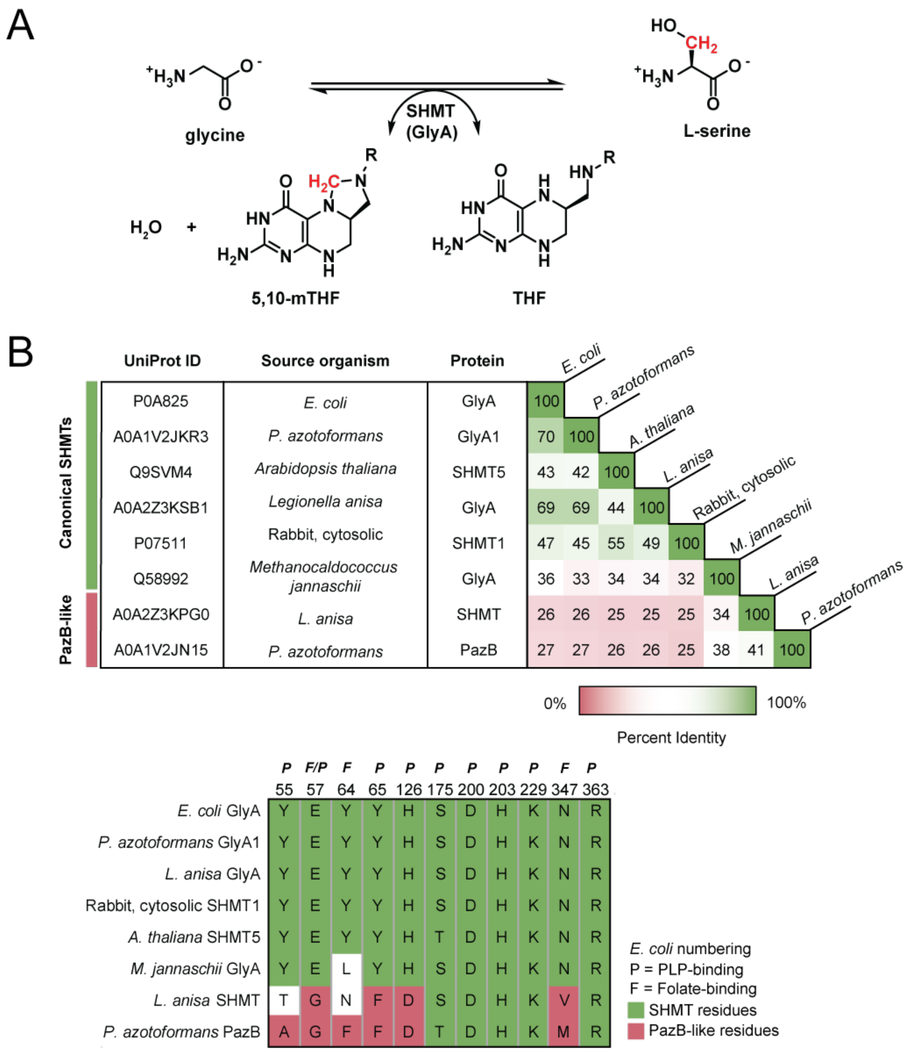

**Figure S3. Gene neighborhoods surrounding the *pazRABC* cluster.** The core *pazRABC* gene cluster is enclosed in a dashed box. The putative gene clusters and their boundaries are identified by gray boxes. The *Pseudomonas* species encode an L-lysine 4-halogenase while the *Legionella* species encode an L-lysine 4,4′-dihalogenase. Some of the genes downstream of the *pazRABC* cluster are labelled, showing that the *pazRABC* cluster is found in a variety of genetic contexts. Interestingly, there are some gene clusters which appear to encode additional enzymes which might be modifying or utilizing pazamine. For example, *P. orientalis* encodes an oxidoreductase and a partner flavodoxin and *L. sainthelensi* could be incorporating a pazamine-like metabolite into a non-ribosomally synthesized peptide.

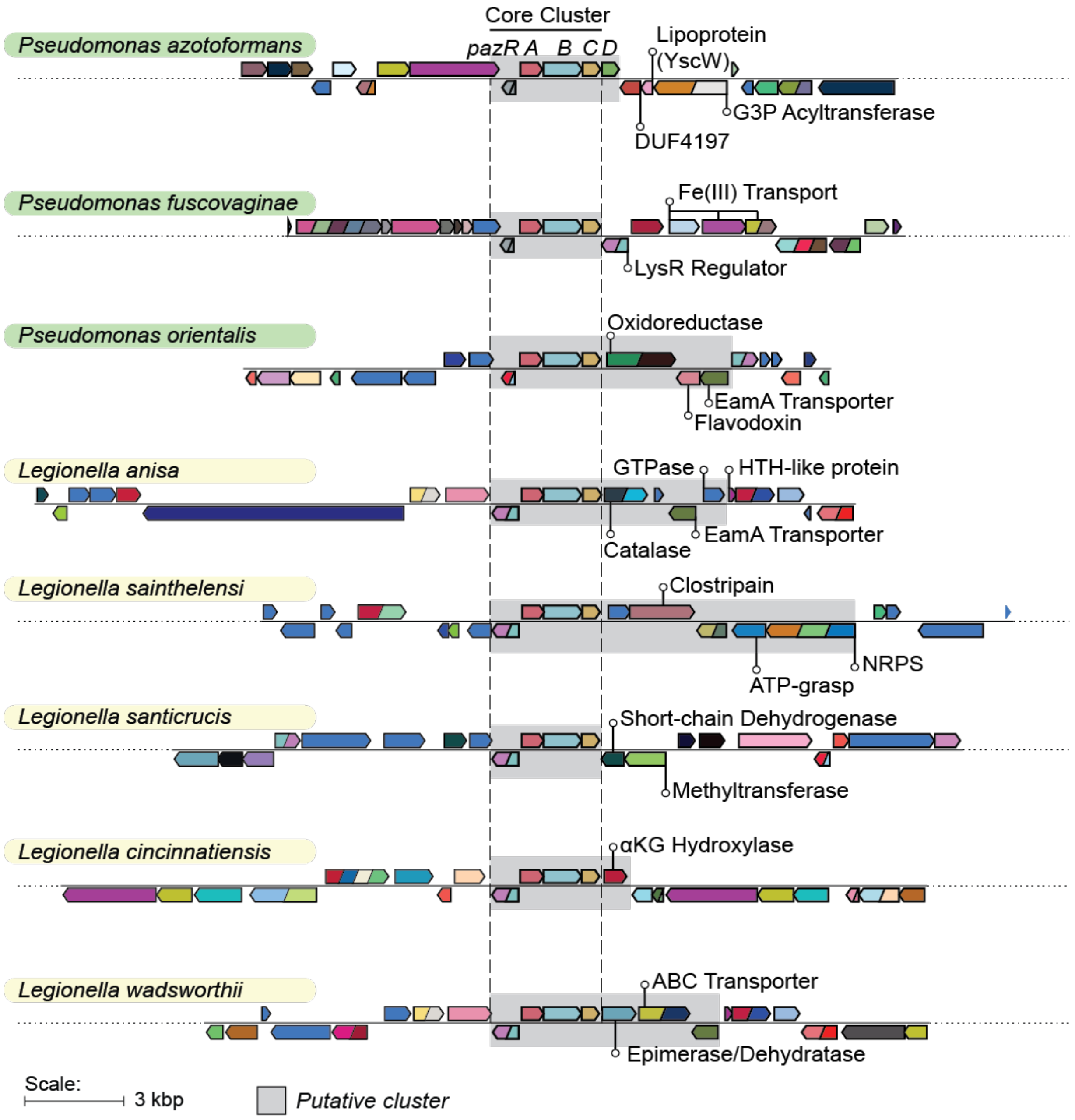

**Table S2. Summary of PazB variants.** A number of PazB variants were cloned for heterologous expression in *E. coli* (unless otherwise noted). While soluble protein was obtained for some variants, activity has not yet been detected *in vitro* for PazB thus far (nd, not detected). Several PazB variant sequences were generated using PROSS [*19*] or ProteinMPNN [*20*], which suggest mutations that will increase solubility (*Table S1*).

| Plasmid(s) | Notes | Solubility | Activity |
|---|---|---|---|
| pMBS3 (pMMPc-PazB-$His_6$) | PazB-$His_6$ in pMMPc for constitutive expression in *P. azotoformans* | nd | nd |
| pMBS8 (pET16hp-$His_{10}$-PazB) | $His_{10}$-PazB in pET16hp | Insoluble | nd |
| pMBS10 (pET16hp-WP_169378548.1) | $His_{10}$-PazB ortholog in pET16hp | Insoluble | nd |
| pMBS11 (pET24a-WP_169378548.1) | PazB-$His_6$ ortholog in pET16hp | Insoluble | nd |
| pMBS15 (pET16hp-PazB_PROSS1) | $His_{10}$-tag, 5% of residues mutated by PROSS, active site preserved | Insoluble | nd |
| pMBS16 (pET16hp-PazB_PROSS3) | $His_{10}$-tag, 8% of residues mutated by PROSS, active site preserved | Soluble | Inactive |
| pMBS17 (pET16hp-PazB_PROSS4) | $His_{10}$-tag, 11% of residues mutated by PROSS, active site preserved | Soluble | Inactive |
| pMBS18 (pET24hp-PazB_PROSS1) | $His_{10}$-tag, 5% of residues mutated by PROSS, active site preserved | Insoluble | nd |
| pMBS19 (pET24hp-PazB_PROSS4) | $His_{10}$-tag, 11% of residues mutated by PROSS, active site preserved | Soluble | Inactive |
| pMBS21 (pET16hp-PaPazB) | $His_{10}$-Npgk-PazB in pET16hp | Insoluble | nd |
| pMBS22 (pET28a-SUMO-PazB) | $His_{14}$-SUMO-PazB in pET28 | Insoluble | nd |
| pMBS24 (pET28a-LaSHMT) | N-terminal $His_6$-tag, PazB ortholog (41% ID) from *Legionella anisa* (A0A2Z3KPG0) | Insoluble | nd |
| pMBS25 (pET28a-LwSHMT) | N-terminal $His_6$-tag, PazB ortholog (39% ID) from *L. wadsworthii* (A0A378LV02) | Insoluble | nd |
| pMBS26 (pET28a-LcSHMT) | N-terminal $His_6$-tag, PazB ortholog (40% ID) from *L. cincinnatiensis* (A0A378IQC9) | Insoluble | nd |
| pMBS33 (pMMPc-StrepPazB) | Strep-PazB in pMMPc for constitutive expression in *P. azotoformans* | nd | nd |
| pMBS39 (pET28a-Ecoli_PazB) | *E. coli* GlyA (P0A825) with active site redesigned to resemble PazB (9 mutations), N-terminal $His_6$-tag | Soluble | Inactive |

| | | | |
|---|---|---|---|
| pMBS40 (pET28a-Salici_PazB) | PazB homolog from *Salicibacter cibi* (JGI IMG: 2994814746) with active site redesigned to resemble PazB (5 mutations), N-terminal $His_6$-tag | Soluble | Inactive |
| pMBS41 (pET21a-m1_21) | ProteinMPNN-designed PazB variant, C-terminal $His_6$-tag | Soluble | Inactive |
| pMBS42 (pET21a-m1_23) | ProteinMPNN-designed PazB variant, C-terminal $His_6$-tag | Soluble | Inactive |
| pMBS43 (pET21a-m2_46) | ProteinMPNN-designed PazB variant, C-terminal $His_6$-tag | Soluble | Inactive |
| pMBS44 (pET21a-m2_48) | ProteinMPNN-designed PazB variant, C-terminal $His_6$-tag | Soluble | Inactive |

**Figure S4. Metabolomic analysis with MS-DIAL.** A metabolomic workflow identified pazamide as a product of the pPazAB cluster. Cultures of *P. azotoformans* containing either the empty pMMPc control plasmid or the pPazAB overexpression plasmid were grown in triplicate, starting from individual colonies, in LB Miller broth supplemented with Gm (30 µg mL$^{-1}$) at 30ºC for 3 d. Methanolic extracts of their metabolomes were analyzed by HILIC/QTOF-MS and the data was processed with MS-DIAL. Shown below is data from the 3-d time point. Three features at the same retention time were identified, all believed to be different adducts of the same metabolite. The exact mass suggests that the higher molecular weight metabolites are sodium adducts of pazamide.

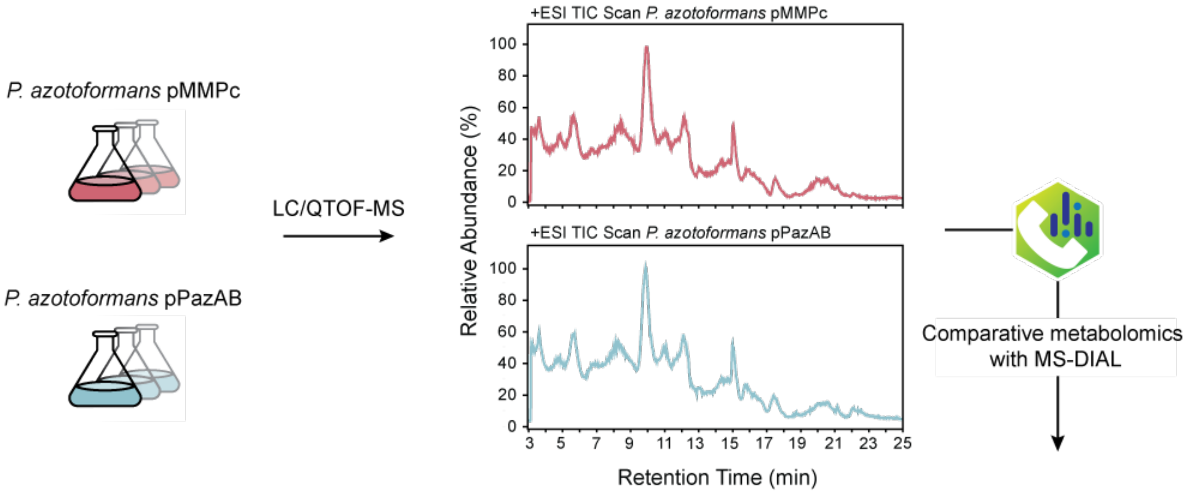


| RT (min) | m/z | Type | ID | ANOVA *p*-value | S/N | Fold-increase |
|---|---|---|---|---|---|---|
| 17.21 | 267.0959 | [M+Na]+ | Pazamide, sodium adduct | 0.167 | 88 | 10300 |
| 17.22 | 289.0785 | [M+2Na-H]+ | Pazamide, disodium adduct | 0.174 | 52 | 6253 |
| 18.74 | 296.1220 | [M+H]+ | Unknown | 0.082 | 15 | 1449 |
| 18.75 | 274.1391 | [M+H]+ | Unknown | 0.093 | 23 | 231 |
| 22.24 | 145.0961 | [M+H]+ | Pazamine | 0.090 | 23 | 179 |
| 17.19 | 245.1117 | [M+H]+ | Pazamide | 0.176 | 135 | 122 |

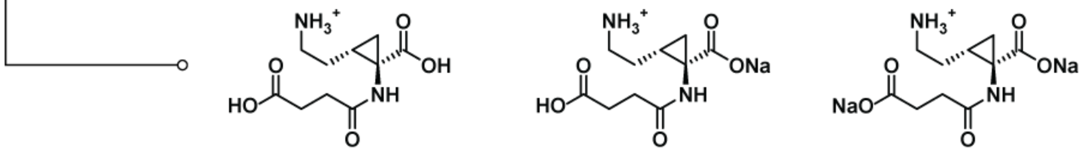

*m/z* = 245.1132 *m/z* = 267.0951 *m/z* = 289.0771

**Figure S5. 1D-NMR characterization of pazamide dimethyl ester (3).** Both spectra are collected at 900 MHz in $D_2O$. (A) $^1H$-NMR. (B) $^{13}C$-NMR.

A

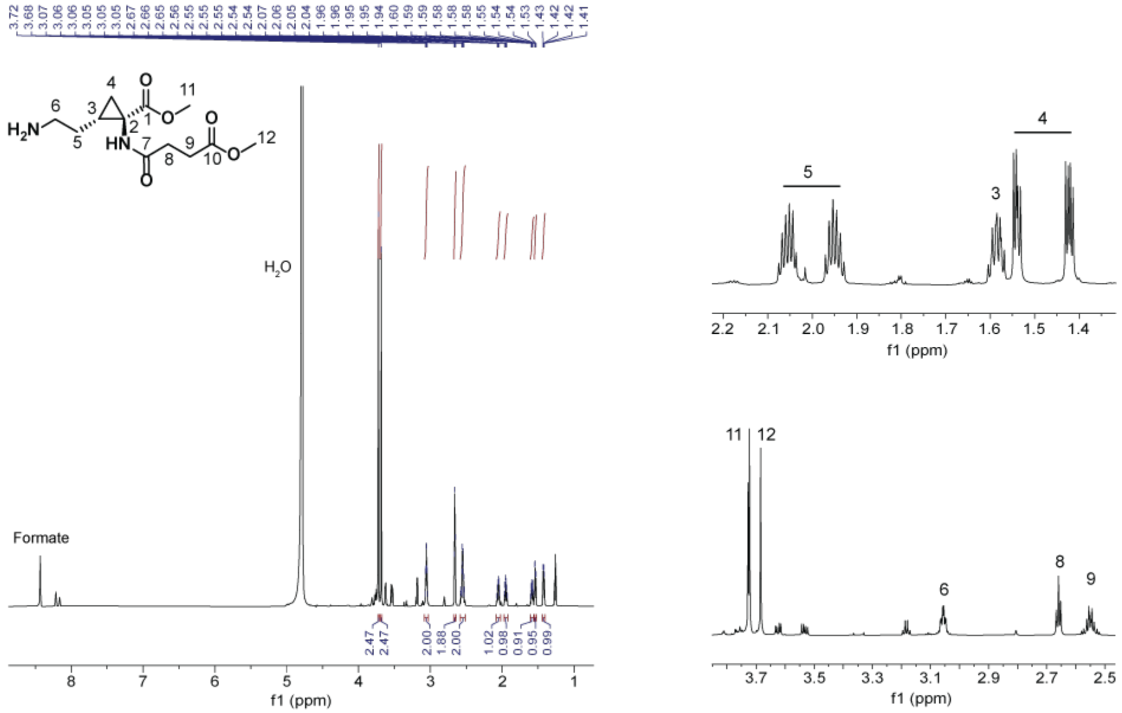

B

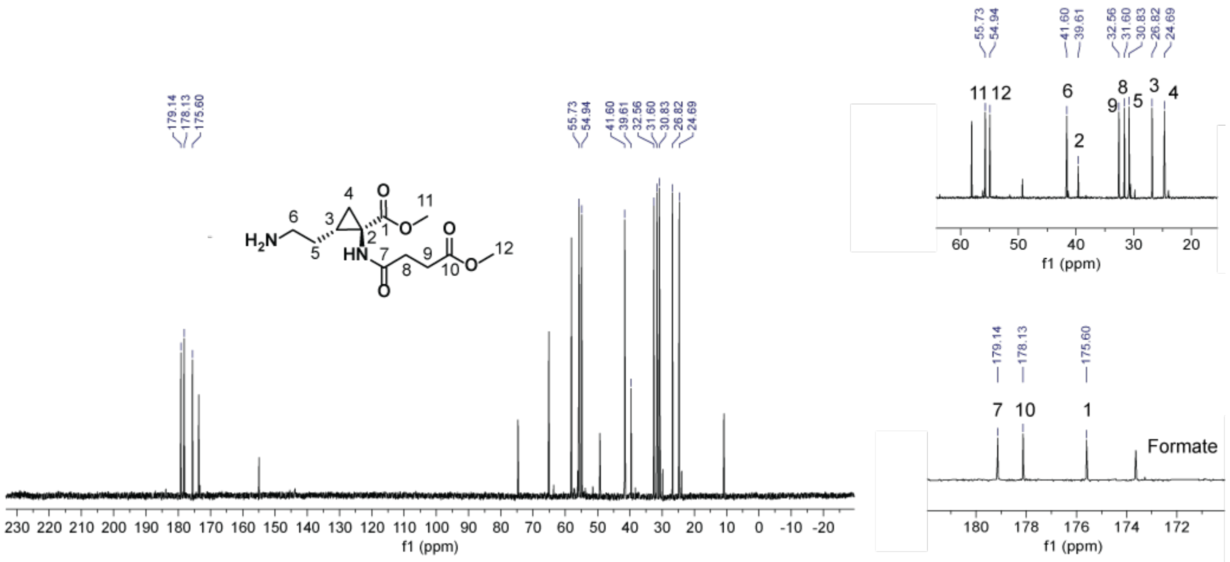

**Figure S6. 2D-NMR characterization of pazamide dimethyl ester (3).** All spectra are collected at 900 MHz in $D_2O$.

(A) Phase-edited HSQC spectrum where the blue phasing represents methylene ($-CH_2-$) units while the red phasing represents methyl ($-CH_3$) and methine (-CH-) carbons. (B) HH-COSY spectrum with COSY correlations indicated with blue squares. Two distinct chains are found which correspond to the succinyl fragment and the amino acid fragment.

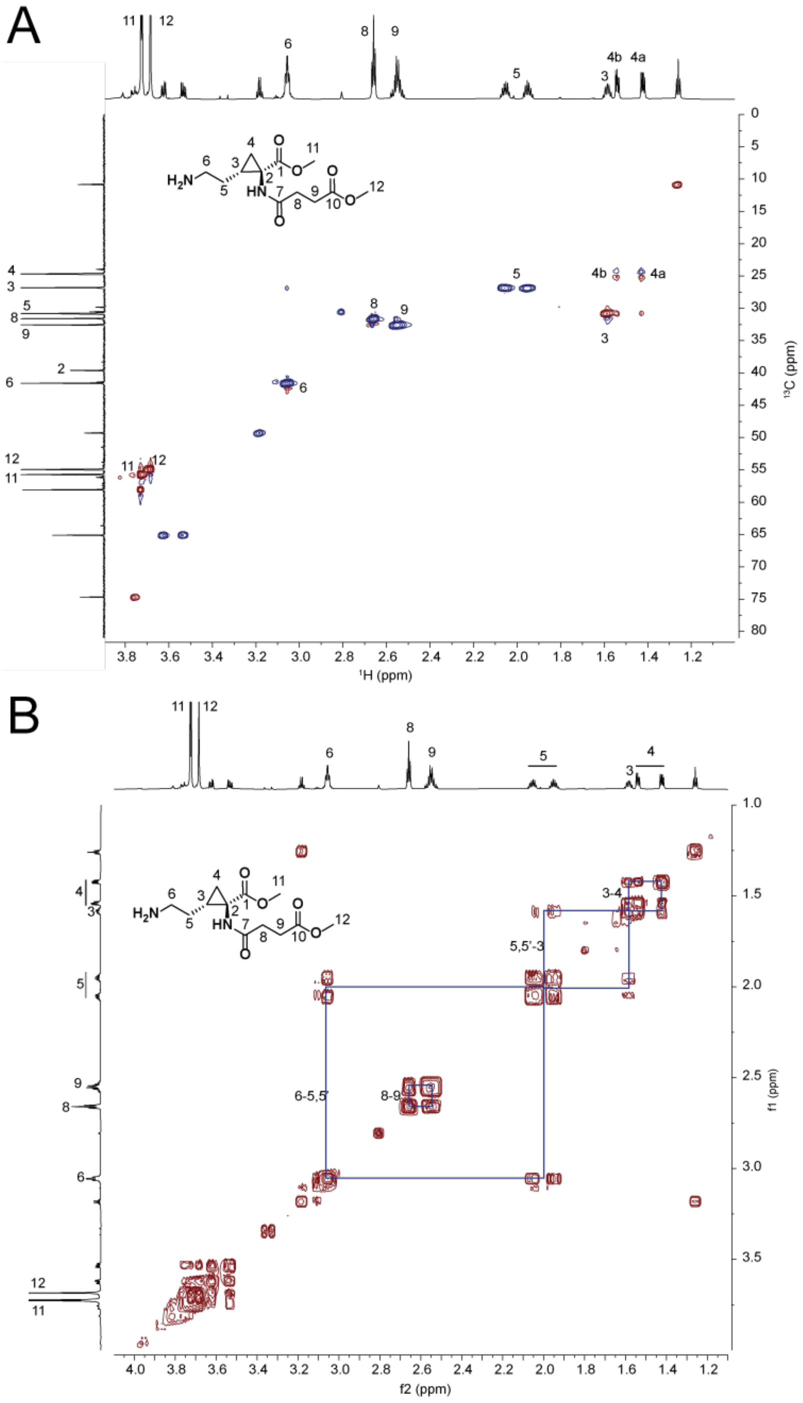

(C) HMBC spectrum with key correlations indicated on the spectrum. Importantly, the tetrasubstituted C-2 was assigned via HMBC correlations to H-5b, H-4a, H-4b, and H-3. This experiment allows us to include the $^{13}$C-NMR signal of C-2 as part of the pazamide structure. Methyl esterification allowed C-1, C-7, and C-10 to be assigned.

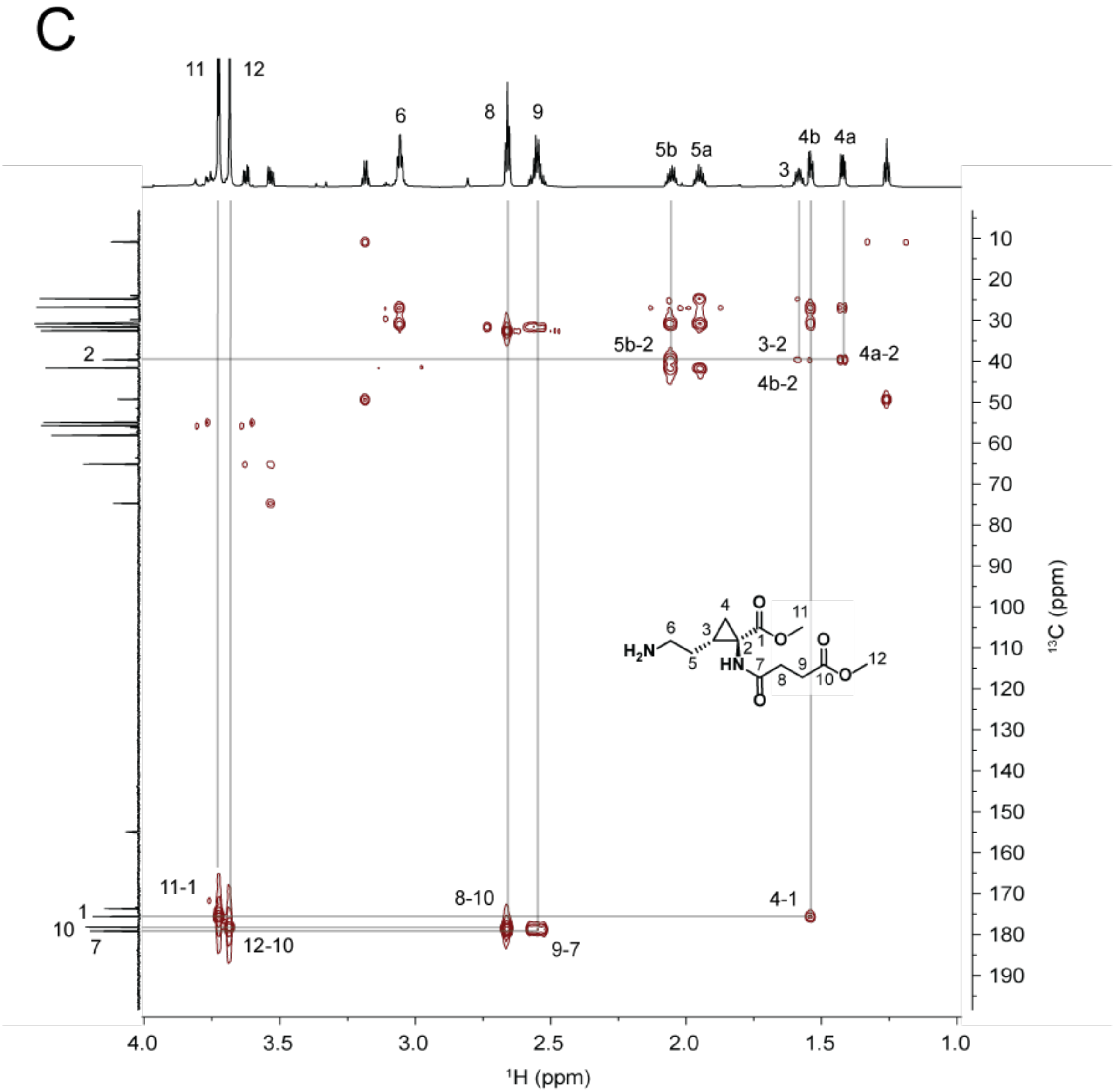

(D) [1]H-NOESY spectra with correlations assigned as labeled. The weak correlation between H-6 and H-11 suggests the aminoethyl moiety and the methyl succinamidyl moiety are on opposite faces of the cyclopropane ring ((*R,R*) or (*S,S*) stereochemistry). (E) Modeling indicates that interaction between H-6 and H-11 would be much weaker or unlikely in the (*S,R*) or the (*R,S*) configurations of **3**. A small population of the (*S,S*) or *(R,R*) isomers is likely to contribute to the observed NOE, with the low barrier (ΔE < 2 kcal/mol) of rotation enabling these enantiomers to access a conformation contributing to the NOE.

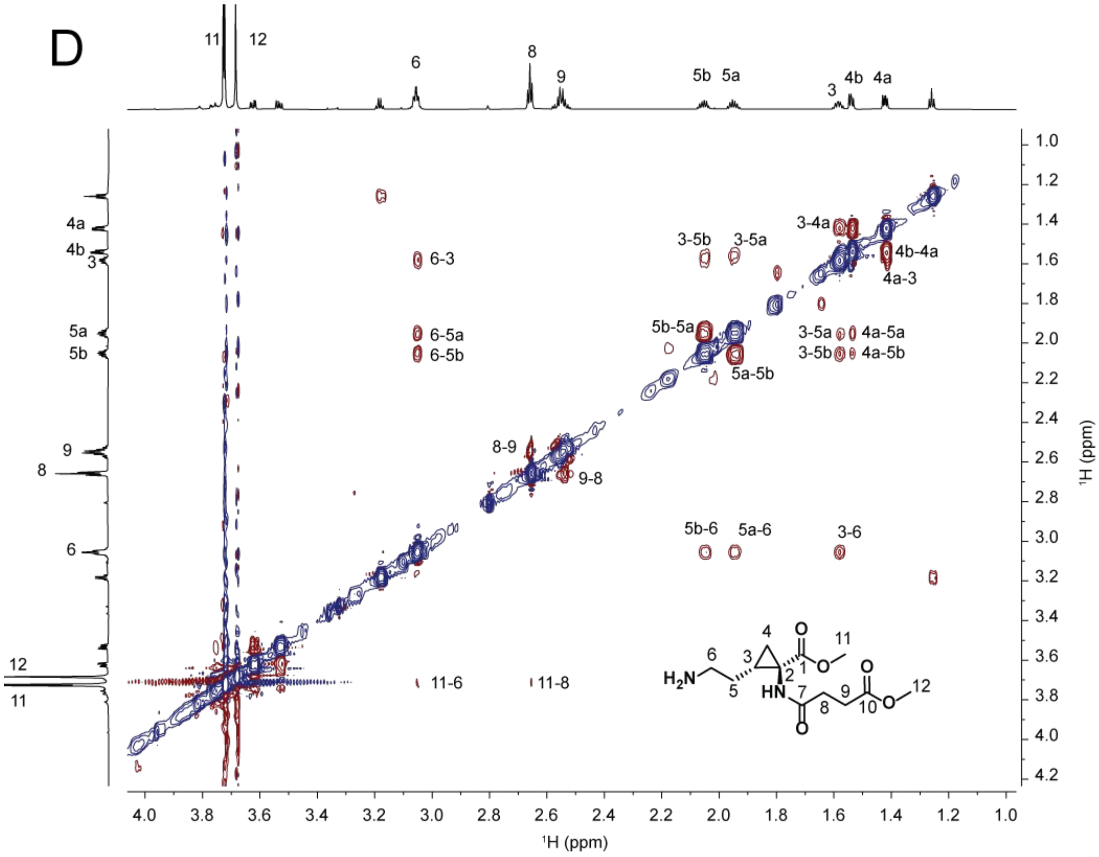


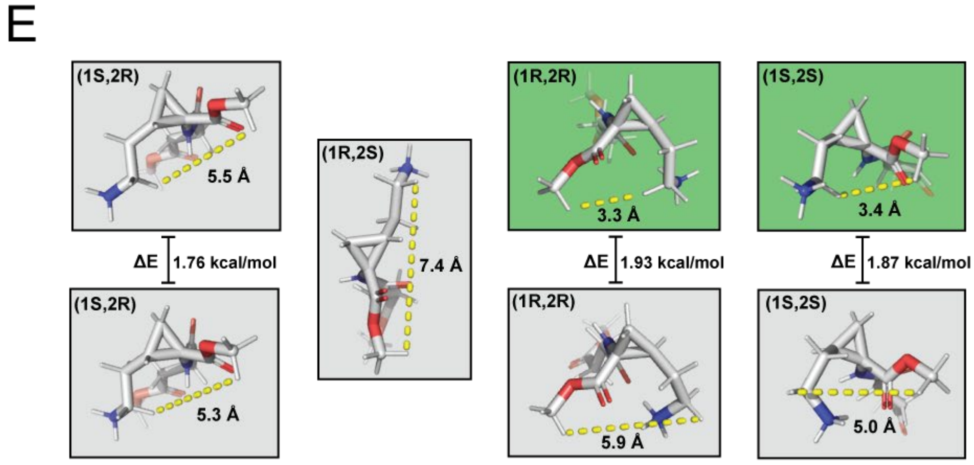

**Figure S7. 1D-NMR characterization of pazamide (2).** (A) $^{1}$H-NMR spectrum of **2** collected at 700 MHz in $D_2O$. (B) $^{13}$C-NMR spectrum of **2** collected at 600 MHz in $D_2O$.

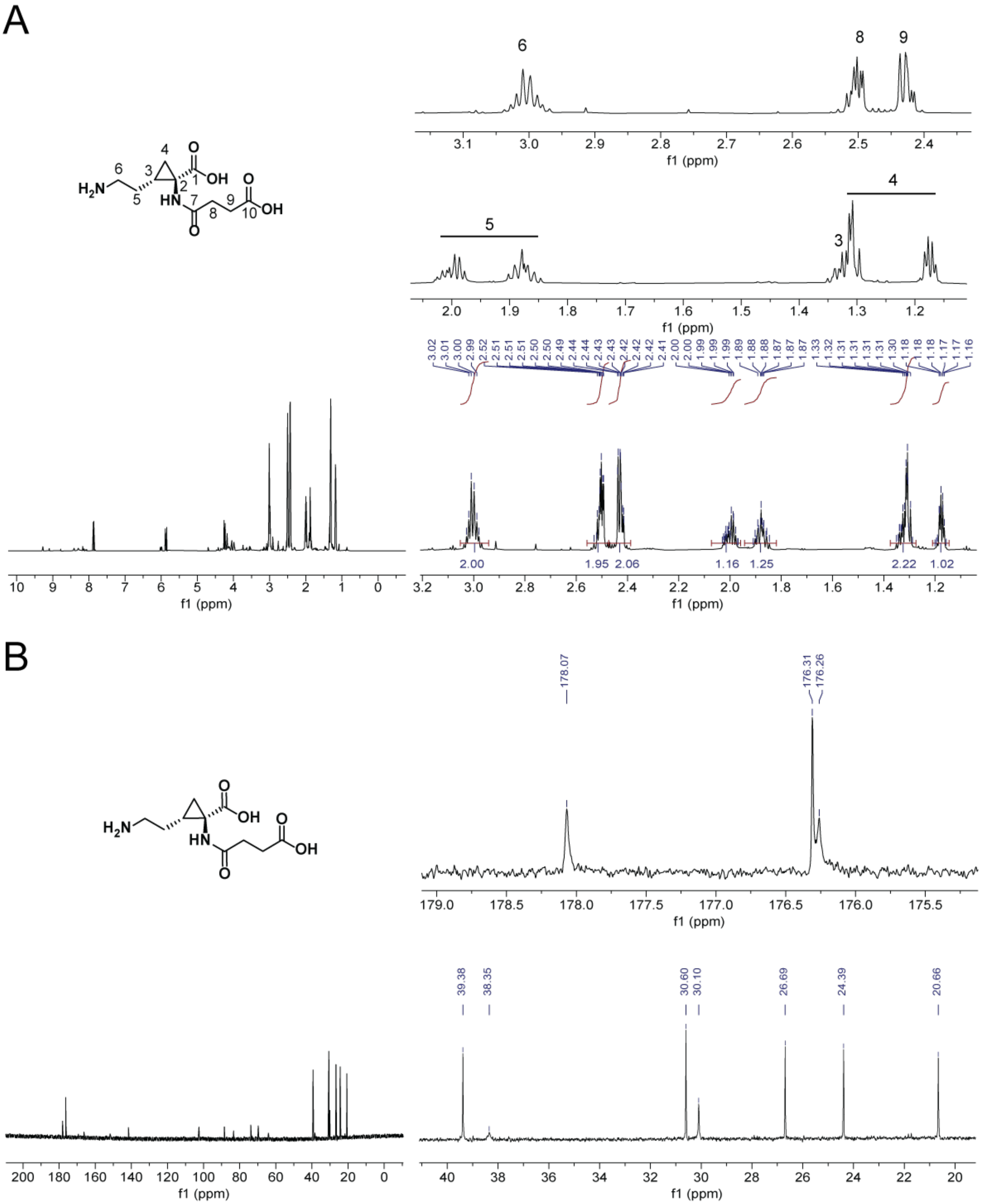

**Figure S8. Confirmation of the PazA product.** (A) The reactions catalyzed by HalA (PfHalA and PazB) and HalB (SiHalB) family members are shown. The stereochemistry of halogenation by PfHalA and SiHalB have previously been reported by our group [*21,22*]. (B) A sequence alignment of PfHalA, PazA, and SiHalB shows a high degree of sequence similarity between PfHalA and PazA (87.8% identity), suggesting that they catalyze chlorination with the same chemical outcome. (C) PazA was heterologously expressed and purified to confirm its activity *in vitro*. LC/MS analysis of the reaction shows that the PazA product has the same retention time as (2*S*,4*R*)-4-chlorolysine produced by PfHalA but a different retention time than the product produced by SiHalB. From these results, PazB is assigned as a L-lysine-4-chlorinase.

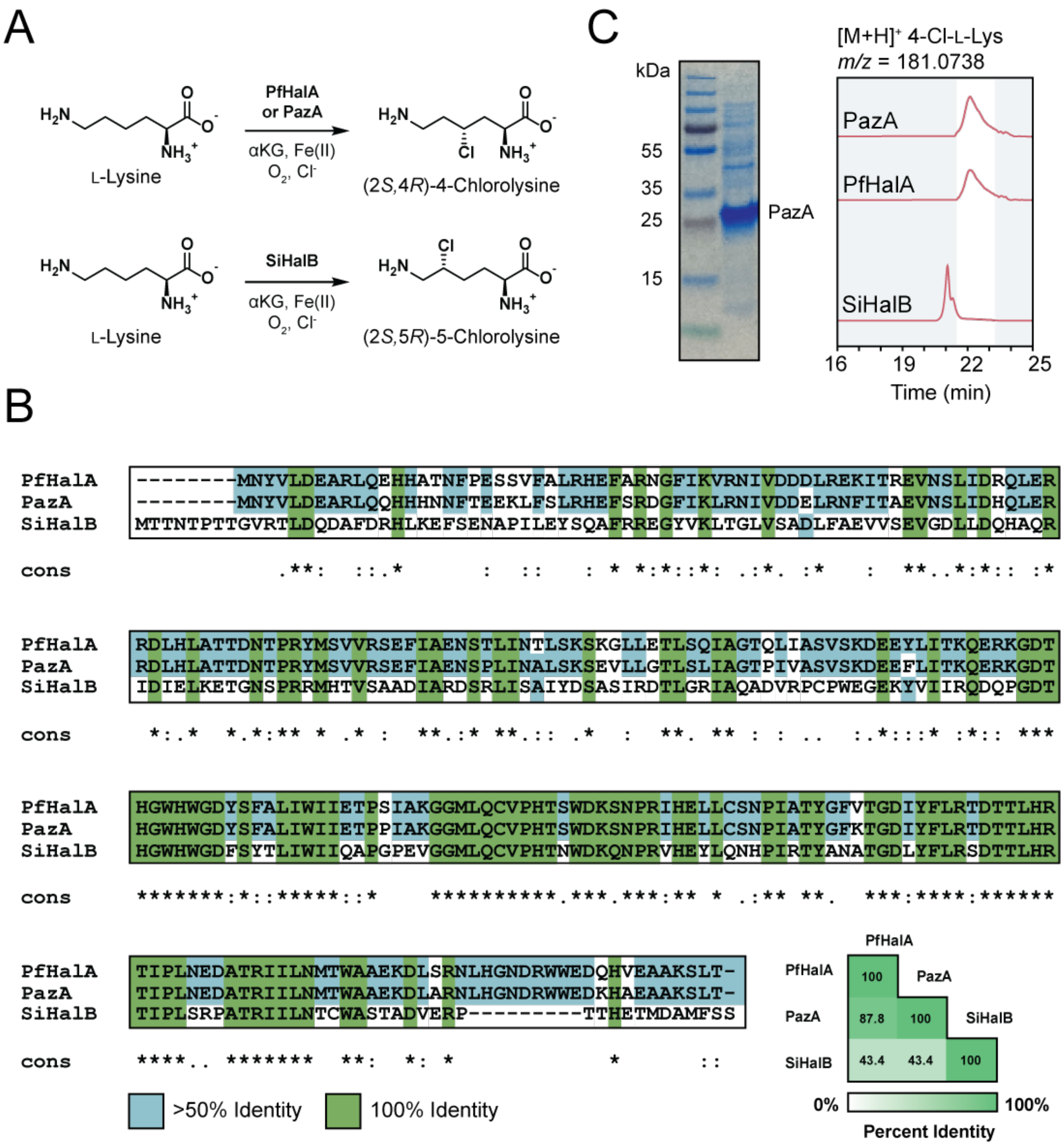

**Figure S9. Structural and sequence analysis of PazB active site.** (A) An AlphaFold model of PazB (teal) overlaid with a crystal structure of *E. coli* GlyA (EcGlyA) (PDB: 1DFO) (red), showing they primarily differ by the presence of an additional N-terminal helix on PazB (dark blue). (B) Comparison of the active sites with a PLP quinonoid intermediate of EcGlyA (glycine) and PazB ((2*S*,4*R*)-4-chlorolysine, **5**). The PazB substrate was docked using AutoDock Vina and computed poses were compared to the glycine-bound crystal structure of EcGlyA to identify a biologically relevant pose. In both the EcGlyA structure and the PazB model, D200/216, H203/219, and R363/R378 are conserved residues which interact with the amino acid-PLP complex through ionic interactions. Both EcGlyA and PazB have two aromatic residues which sit above the active site, Y64/F79 and Y65/F80. While EcGlyA Y64 is known to π-stack with the *p*-aminobenzoic acid ring of the folate cofactor [*13*], PazB F79 is predicted to be in a rotated conformation relative to EcGlyA Y64, potentially to help fold 5-PLP into a reactive conformation. Additionally, mutational analysis of EcGlyA Y65 to Y65F has suggested that the phenolic hydroxyl group is important for weaking the strength of the ionic interaction between the substrate carboxylate and R363, as well as promoting substrate release [*23*]. The analogous position in PazB, F80, is natively phenylalanine. In EcGlyA, H126 π-stacks with the PLP pyridinium ring. This is consistent with the fact the fold type I PLP-dependent enzymes (which includes SHMT) have a π-stacking aromatic residue at this position [*18*]. Conversely, this residue is an aspartate in PazB, D141. A polar pocket composed of D141, T142, and S143 appear poised to interact with the terminal ammonium of the substrate, potentially aiding 5-external aldimine in folding into a reactive conformation.

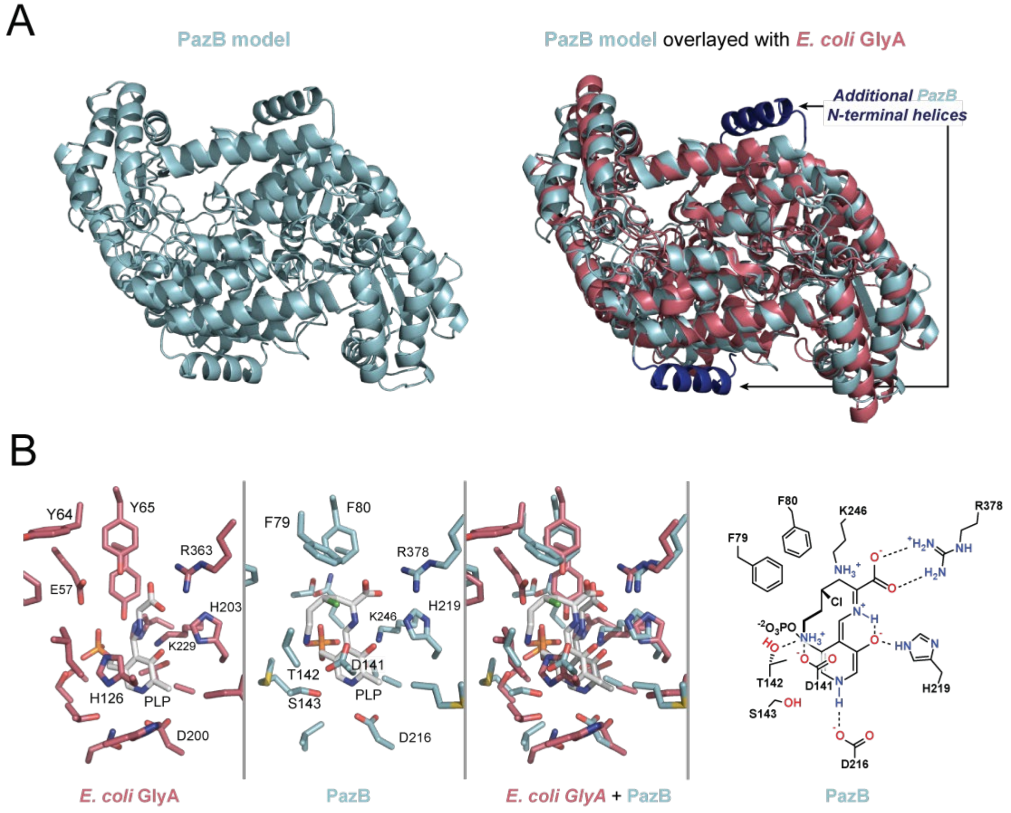

**Figure S10. Arginine/ornithine succinyltransferase can succinylate pazamine.** (A) Scheme of an abbreviated arginine utilization pathway. Arginine and ornithine, a metabolic precursor to arginine, can both be succinylated at $N_\alpha$ by the enzyme AOST, encoded by the genes *aruF* and *aruG* [*24*]. Succinylation is the entry point into the degradation of L-Orn and L-Arg into L-Glu and succinate. (B) As **1** is structurally similar to L-Orn we hypothesized that AOST could succinylate the $N_\alpha$ of **1**. (C) Chromatograms for **1** (*m/z* = 145.0972 $[M+H]^+$) and **2** (*m/z* = 245.1132 $[M+H]^+$) production are shown. While *P. azotoformans* pPazAB can produce both **1** and **2**, the AOST-deficient strain *Pseudomonas azotoformans* Δ*aruFG* pPazAB only biosynthesizes **1**. The parent strain containing the empty plasmid pMMPc does not produce **1** or **2** under these conditions.

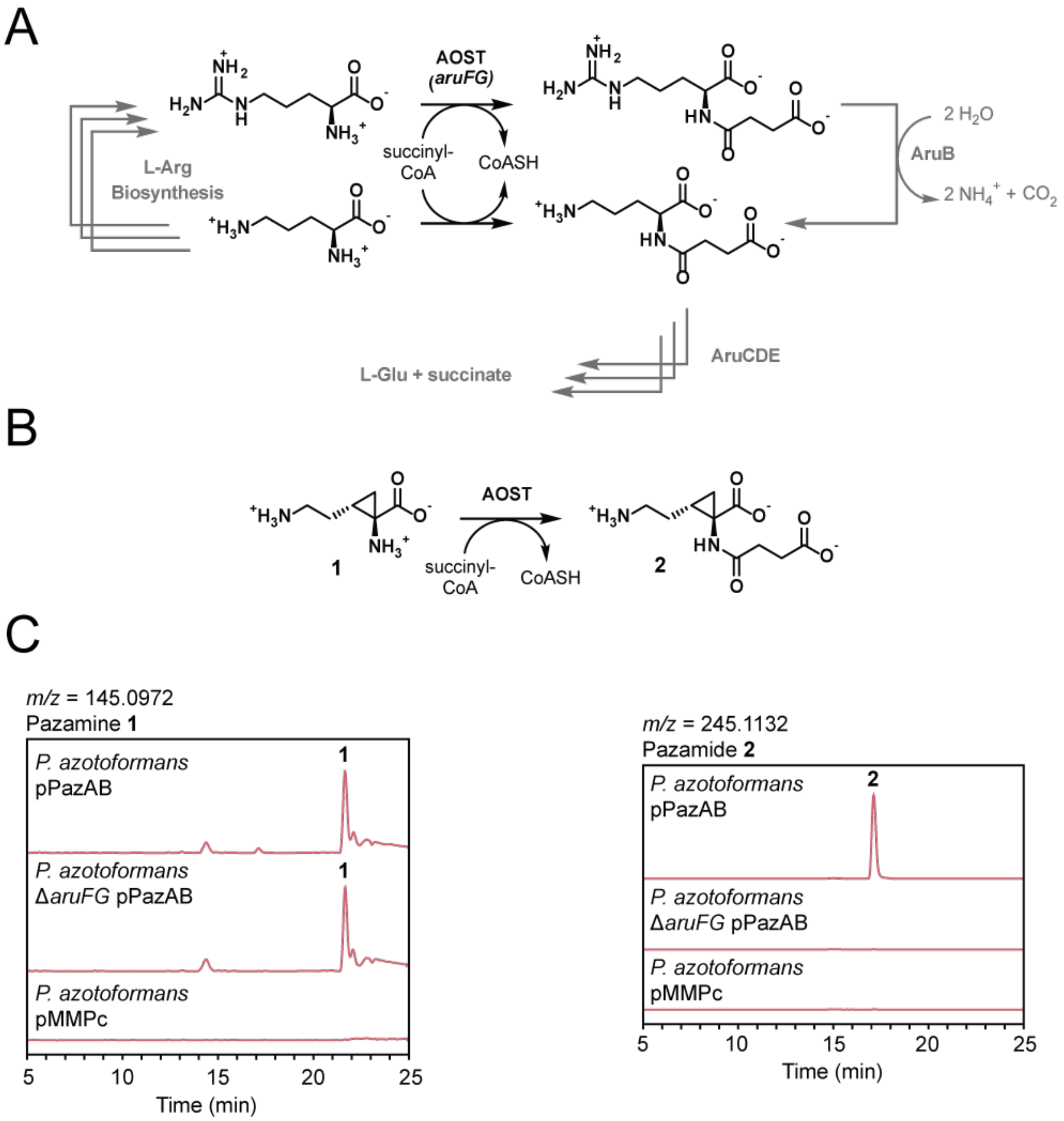

**Figure S11. *Arabidopsis* seedling growth phenotype after bacterial inoculation.** (A) The quantification of the total root lengths of inoculated seedlings over 14 d. Bars and error bars denote the mean ± s.d., and the line plot denotes change over time. The numbers indicate statistically different means (ANOVA and Tukey post-hoc test) between groupings at individual time points, described below the graph (*n*: NBI (no bacterial inoculation) = 18, ΔAOST = 19, pPazAB = 20, ΔpazA = 25, and WT = 22). (B) Photos of *A. thaliana* at 14 d post inoculation display the differences in root phenotypes linked to bacterial inoculation by different *P. azotoformans* strains.

A

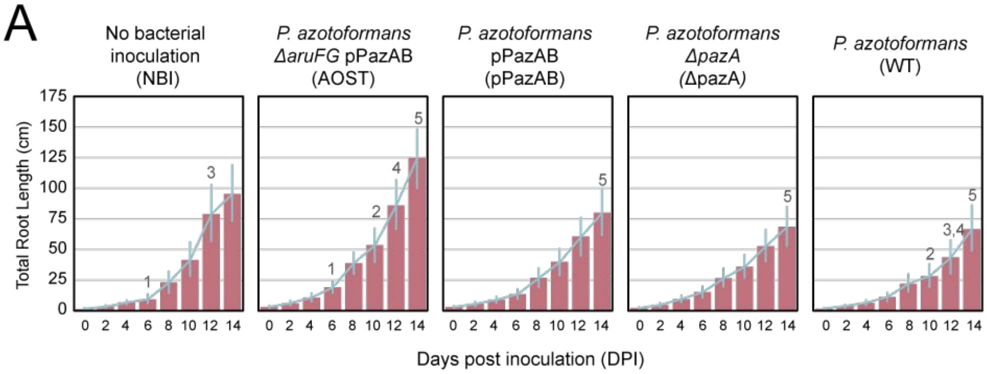


| Annotation | Group 1 | Group 2 | *p*-value |
|---|---|---|---|
| 1 | ΔAOST | NBI | 0.0374 |
| 2 | ΔAOST | WT | 0.0255 |
| 3 | NBI | WT | 0.0499 |
| 4 | ΔAOST | WT | 0.0087 |
| 5 | ΔAOST | pPazAB | 0.0378 |
| 5 | ΔAOST | ΔpazA | 0.0021 |
| 5 | ΔAOST | WT | 0.002 |

B

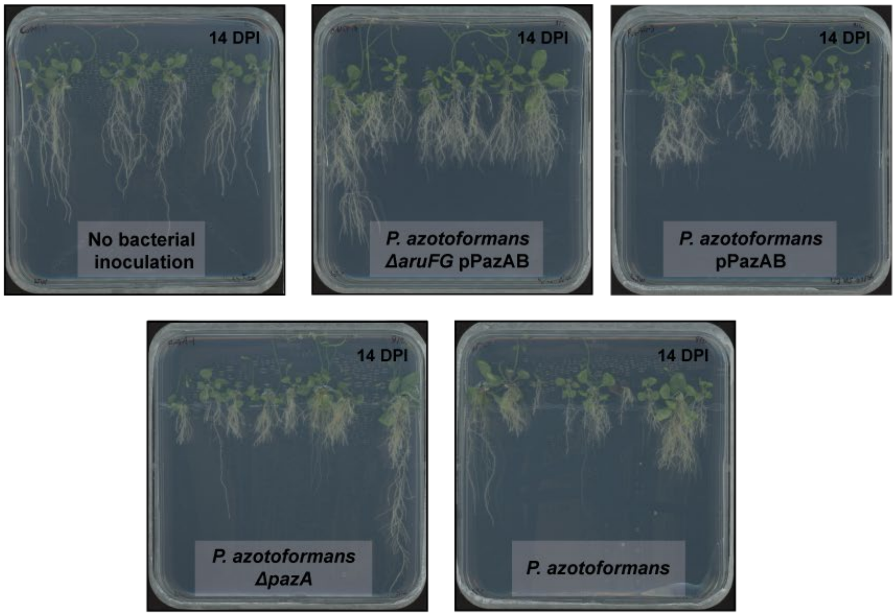


*Arabidopsis thaliana*

**Figure S12. Deuterium labeling supports PazB-mediated cyclization.** (A) 4- and 5-L-chlorolysine can either be cyclized by PazB to their corresponding 3- or 4-membered carbocycle or they can degrade by intramolecular lactonization [*25*]. Using *d*9-L-lysine enables us to distinguish the two products by their unique *m/z* values. (B) A metabolite with *m/z* ($[M+H]^+$) = 152.411 is detected when *P. azotoformans ΔaruFG* pPazAB is fed *d*9-L-lysine and assigned as *d*7-**1**. (C) Following Fmoc-derivatization, $C_{18}$ reverse-phase LC/QQQ-MS experiment is used to detect Fmoc-derivatized amino acid products. The chromatograms demonstrate detection of Fmoc-*d*7-**1** (left) and Fmoc-*d*7-**7** (right).

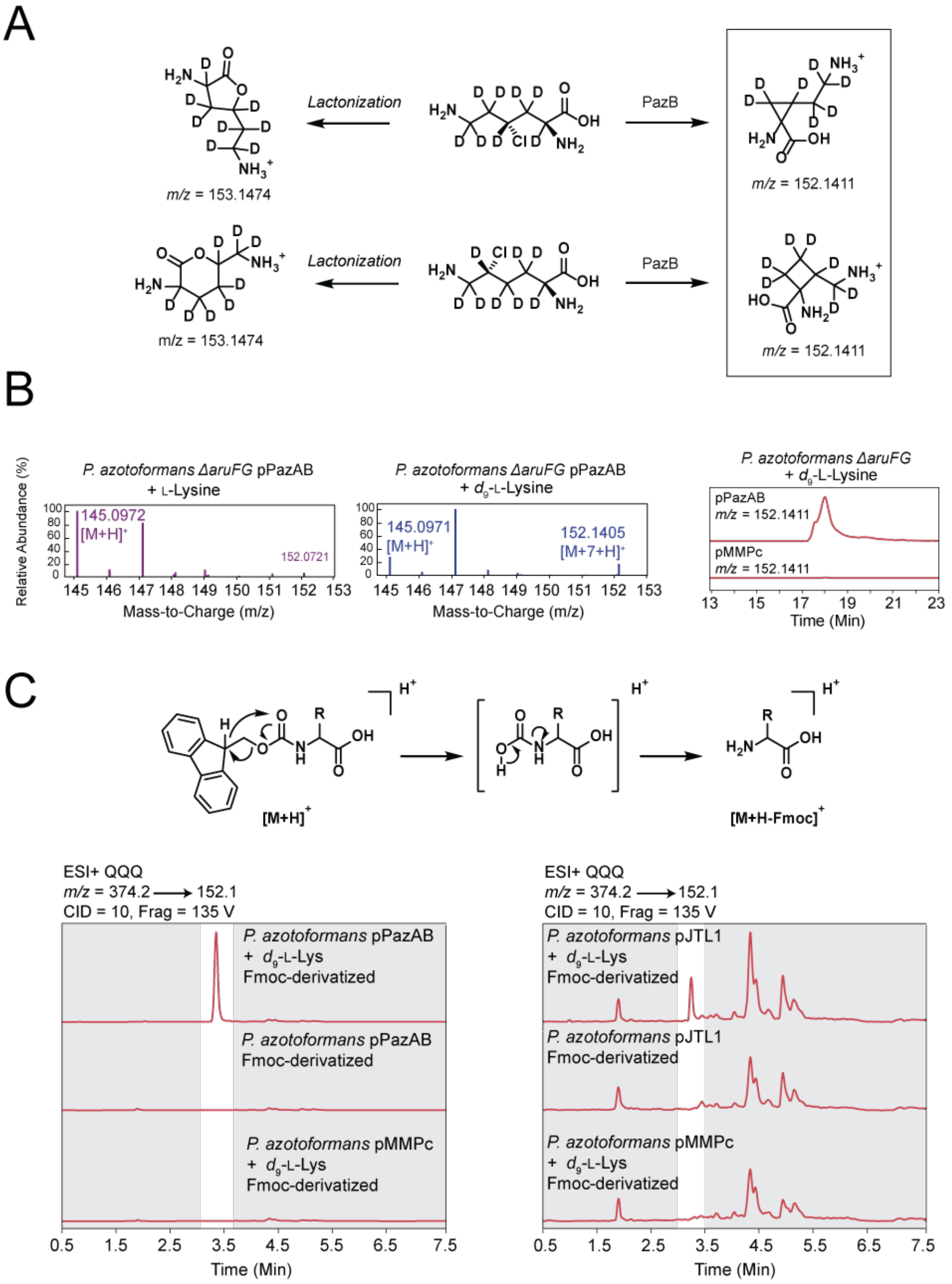

**Author Contributions**. M. B. Sosa was responsible for designing experiments, performing experiments, collecting data, analyzing data, and writing the manuscript. J. T. Leeman was responsible for designing experiments, performing experiments, and collecting data. L. J. Washington was responsible for designing experiments, performing experiments, collecting data, and analyzing data. H. V. Scheller was responsible for designing experiments and validating data. M. C. Y. Chang was responsible for administering the project, designing experiments, validating data, and writing the manuscript. All authors were involved in editing the manuscript.
